# Coordinate regulation of Activin and Bone Morphogenetic Protein signaling by a lysosomal trafficking switch

**DOI:** 10.1101/2024.01.29.577837

**Authors:** Warren W. Hom, Senem Aykul, Lampros Panagis, Krunal Patel, Susannah Brydges, Erich J. Goebel, Kaitlin N. Lydon, John B. Lees-Shepard, Sarah J. Hatsell, Vincent Idone, Aris N. Economides

**Affiliations:** Regeneron Pharmaceuticals, 777 Old Saw Mill River Road, Tarrytown, NY 10591

## Abstract

BMP/TGFß family ligands act mainly as factors that differentially initiate Smad1/5/8 or Smad2/3 signaling via heterotetrameric complexes comprised of two type I and two type II receptors (IIR). ActA (ActA) stands out as it activates Smad2/3 signaling through type I receptor ACVR1B, whereas it generates non-signaling complexes (NSCs) with ACVR1. In fibrodysplasia ossificans progressiva (FOP), a genetic disorder caused by missense mutations in ACVR1 (ACVR1.FOP), ACVR1.FOP•ActA•IIR complexes activate Smad1/5 signaling, mimicking those formed with BMPs. As the NSCs that ActA forms with ACVR1 are stoichiometrically identical with signaling complexes formed with ACVR1.FOP, we explored how NSCs differ from their signaling counterparts. We show that ACVR1•ActA•IIR complexes rapidly traffic to lysosomes, where their constituent components are degraded, reducing the cell’s responsiveness to BMPs along with ActA’s availability. This property is specific to ActA as Activin B, AB, and AC do not form lysosomal trafficking complexes with ACVR1, but rather remain on the surface as NSCs. Lysosomal trafficking and degradation of ACVR1•ActA•IIR complexes is a novel regulatory mechanism of BMP/TGFß signaling whose physiological roles remain largely unexplored.

## Main

BMPs and TGFßs have primarily been studied as factors that initiate Smad signaling through heterotetrameric complexes of their corresponding type I and type II receptors ^1–4^. BMP3, lefty1, lefty2, Inhibin A, and Inhibin B stand as exceptions as they function solely as antagonists. This is accomplished by forming non-productive complexes with type II receptors, either alone ^5, 6^ or in combination with non-signaling coreceptors ^7, 8^, thereby rendering type II receptors unavailable for signaling.

Recently, a more complex picture emerged with the Activin subfamily during exploration of the molecular mechanisms that drive the pathophysiology of fibrodysplasia ossificans progressiva (FOP). FOP is an autosomal-dominant genetic disorder characterized by progressive and cumulative heterotopic ossification (HO) of connective tissues and arising from heterozygous missense mutations in the region encoding the cytoplasmic domain of the type I BMP receptor ACVR1 ^9^. The great majority of patients carry a variant altering Arginine 206 to Histidine (*ACVR1^R206H^*) ^10^. This amino acid change renders ACVR1.R206H responsive to ActA(ActA) as well as Activin B, AB, and AC ^11, 12^, a property shared with all other FOP-causing variants of ACVR1 (ACVR1.FOP) ^12^. Importantly, inhibition of ActA using monoclonal antibodies ameliorated HO in FOP mice (*Acvr1^R206H/+^*) ^11, 13, 14^, resulting in exploration of ActA as a drug target for FOP ^15, 16^.

From a molecular mechanism standpoint, this finding indicated that ActA must interact with ACVR1, as had been initially described ^17–19^. We discovered that ActA brings together preformed heterodimers of ACVR1 and associated type II receptors (IIR) to form ACVR1•ActA•IIR non-signaling complexes (NSCs) ^4, 11,20^. These NSCs inhibit ACVR1-mediated BMP signaling ^11^. However, efficient inhibition is not driven solely by occupancy of type II receptors by ActA as had been previously suggested ^21, 22^ – rather it requires interaction of ActA with ACVR1 ^20^. In addition to interacting with ACVR1, ActA associates with a different type I receptor, ACVR1B ^23–25^ and activates Smad2/3 signaling ^26, 27^. Hence, in contrast to BMP3, lefty 1, lefty 2, Inhibin A, and Inhibin B, which act solely as inhibitors of signaling, ActA has a dual role: it inhibits BMP-induced Smad1/5 signaling mediated by ACVR1, whereas it activates Smad2/3 signaling from ACVR1B.

In our initial attempts to explore the physiological role of **ACVR1•ActA•IIR** NSCs, we engineered an ActA mutein, ActA.Nod.F2TL (F2TL), that retains the ability to activate ACVR1B but cannot form NSCs with ACVR1 ^20^. Although F2TL cannot form NSCs with ACVR1 (indicating lack of binding), it is nonetheless able to activate Smad1/5 signaling via ACVR1.R206H. The reason for this is two-fold:

ACVR1 exists in ligand-independent heterodimeric complexes with partner type II receptors, and ACVR1.R206H can be activated by simple dimerization irrespective of the dimerizing agent ^4^. Therefore, F2TL can still dimerize **ACVR1.R206H•IIR** heterodimers by binding the type II receptors, precipitating the formation of heterotetramers comprised of two ACVR1.R206H•IIR dimers even in the absence of a direct interaction of F2TL with ACVR1. We took advantage of this property to explore the role of NSCs in FOP, by comparing the amount of heterotopic bone that forms in response to equivalent amounts of ActAversus F2TL in FOP mice. As FOP is a heterozygous condition, we hypothesized that F2TL by escaping engagement into NSCs (that would be formed with ACVR1 expressed from the remaining wild type allele of *Acvr1*) and generate more heterotopic bone in an implant assay. FOP mice with F2TL implants generated approximately 6 times more heterotopic bone at the implant site compared with FOP mice receiving identical implants containing an equimolar amount of ActA^20^. These data indicated that in FOP mice **ACVR1•ActA•IIR** NSCs play an important role in that they temper the amount of heterotopic bone formed presumably by reducing the levels of ActA and consequently the level of activation of ACVR1.R206H ^20, 28^. However, these findings did not address the molecular mechanism by which NSCs bring about their effects. To address this question, we explored the fate of **ACVR1•ActA•IIR** NSCs at the cellular level as well as and how NSCs affect signaling.

## Results

### ActA-ACVR1 NSCs are endocytosed and traffic to lysosomes

To assess the fate of **ACVR1•ActA•IIR** NSCs we generated versions of ACVR1 and ActA that can be tracked using fluorescence microscopy. For ACVR1, we engineered fusion proteins, Halo-ACVR1-HiBiT and Halo-ACVR1.R206H-HiBiT, which are N-terminally tagged with Halo tag and C-terminally tagged with HiBiT tag. This enables tracking of Halo-ACVR1-HiBiT and Halo-ACVR1.R206H-HiBiT using the fluorescent ligand (AF660) which emits in the red spectrum. To avoid artifacts associated with overexpression ^4^ and to reduce competition from untagged ACVR1, we introduced these ACVR1 fusion proteins into *Acvr1* homozygous-null (*Acvr1^−/-^*) mouse embryonic stem cells (mESCs), using lentiviral delivery. The fusion proteins expressed in mESCs (**Extended Data Fig. 1a**), localized on the cell surface (**Extended Data Fig. 1b, c**), and displayed the expected responsiveness to ActA and BMP7 (**Extended Data Fig. 2a, b**). The level of expression of endogenous type II receptors as well as other type I receptors were unaffected by the expression of Halo-ACVR1-HiBiT and Halo-ACVR1.R206H-HiBiT (**Extended Data Fig. 3a**). ActA and F2TL were rendered fluorescent by crosslinking fluorophore AF555 that emits in the yellow spectrum. These fluorescently labeled ligands activated signaling at levels equivalent to their unmodified counterparts, indicating that the fluorophore had not significantly altered interactions with their receptors (**Extended Data Fig. 4**).

Having validated these cell lines and ligands, we used them to explore the fate of different complexes involving ActA. When *Acvr1^−/-^; Tg[Halo-Acvr1-HiBiT]* mESCs are treated with ActA-AF555, Halo-ACVR1-HiBiT along with ActA-AF555, localize in punctate structures within the cytosol (**Fig. 1a; Extended Data Fig. 5a**). In contrast, when *Acvr1^−/-^; Tg[Halo-Acvr1.R206H-HiBiT]* mESCs are treated with ActA-AF555, neither Halo-ACVR1.R206H-HiBiT nor ActA are found in the cytosol at appreciable levels (**Fig. 1a; Extended Data Fig. 5b**). Endocytosis of NSCs that form between Halo-ACVR1-HiBiT and ActA-AF555 proceeds rapidly, being complete approximately 15 minutes after addition of ligand (**Fig. 1b**). Endocytosed NSCs traffic to the lysosome as they colocalize with LAMP1, a lysosomal marker (**Fig. 1a**). These results provide the first evidence that **ACVR1•ActA•IIR** NSCs undergo endocytosis and traffic to lysosomes whereas their ACVR1.R206H counterparts (which signal in response to ActA) do not.

**Figure 1.**
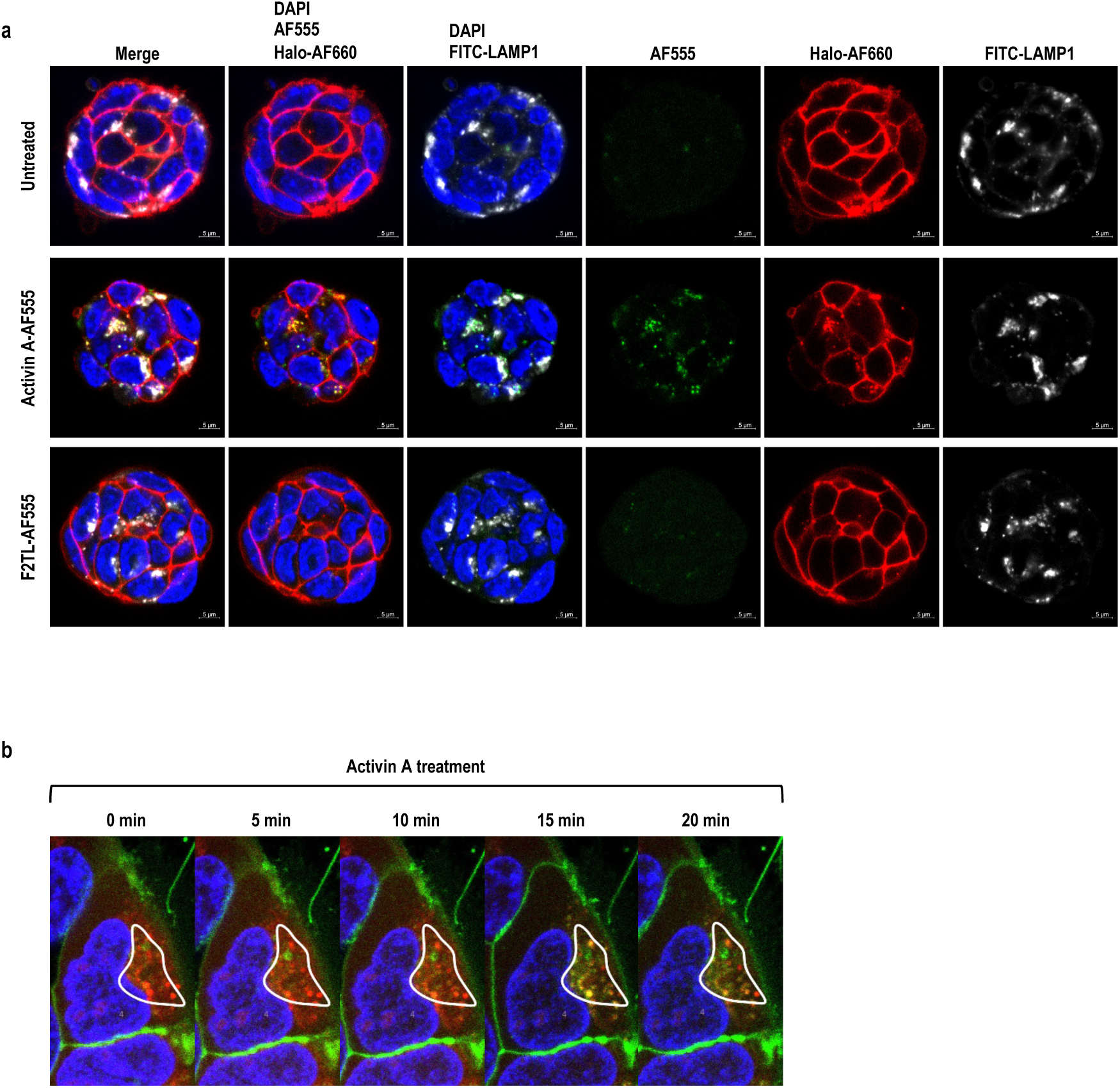
ACVR1•ActA•IIR NSCs traffic to lysosomes. **a**, *Acvr1^−/-^* mESCs expressing Halo-ACVR1-HiBiT were treated with 30nM ActA-AF555 (green) or left untreated. ACVR1 proteins were visualized using AF660 Halo ligand (red). Lysosomes and nuclei were visualized with FITC-LAMP1 (white) and DAPI (blue), respectively. Confocal microscopy was used for imaging. ActA-AF555 co-internalized with Halo-ACVR1-HiBiT and localized in LAMP1-positive lysosomes. In contrast, F2TL-AF555 did not internalize in mESCs expressing Halo-ACVR1-HiBiT. **b**, Live cell imaging of HEK293 cells overexpressing ACVR1-GFP (green) was performed post-lysosome staining with lysopainter (red). The GFP signal was bleached in the outlined area (white), followed by the addition of 30nM ActA. Trafficking of ACVR1-GFP into lysosomes was monitored over a 30-minute period. Images from selected time points are displayed. ACVR1-GFP trafficked to the lysosome within 15 minutes after ActA treatment.

To make certain that endocytosis of **ACVR1•ActA•IIR** NSCs is not a property restricted to mESCs, we tested whether ActA-AF555 also internalizes in the osteoblast-like cell line W20 ^29^ and compared it to F2TL-AF555. In W20 cells, ActA-AF555 localizes in punctate structures within the cytosol whereas F2TL-AF555 is not internalized (**Extended Data Fig. 6)**, mirroring the results obtained in mESCs. These results indicate that endocytosis of **ACVR1•ActA•IIR** NSCs is not a property restricted to one type of cell.

### Lysosomal degradation of NSCs reduces responsiveness to BMPs

Since **ACVR1•ActA•IIR** NSCs traffic to lysosomes, we tested whether the components of the complex are degraded. In HEK293 cells coexpressing myc-tagged ACVR1 (myc-ACVR1) and HA-tagged ACVR2B (HA-ACVR2B), addition of ActA results in reduction of the majority of both myc-ACVR1 and HA-ACVR2B (**Fig. 2a)**. In contrast, with myc-ACVR1.R206H, the levels of myc-ACVR1.R206H and HA-ACVR2B are not reduced after exposure to ActA. This data is consistent with differential trafficking of the corresponding complexes where complexes with wild type ACVR1 traffic to lysosomes whereas complexes with ACVR1.R206H do not.

**Figure 2.**
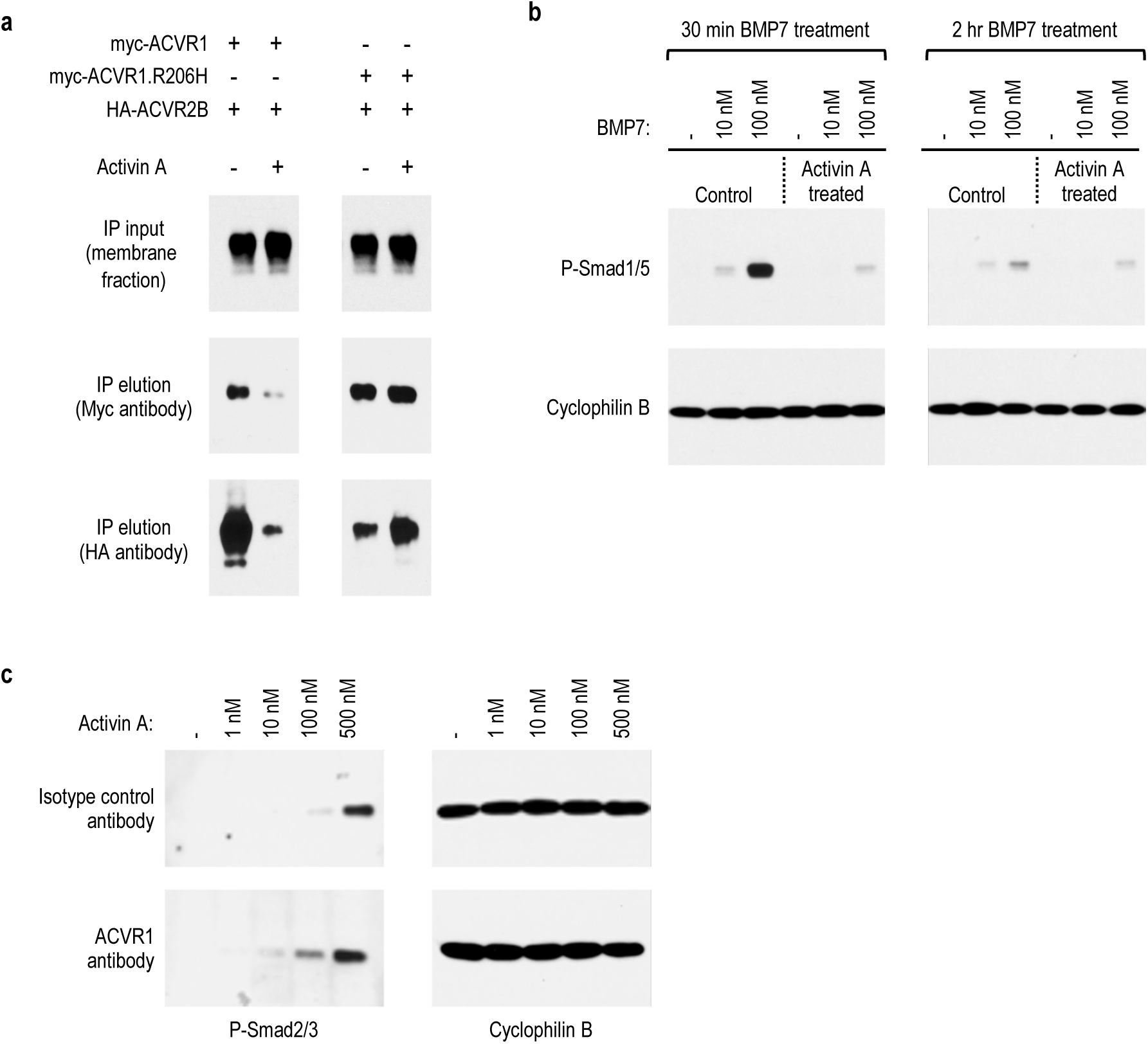
ACVR1•ActA•IIR NSCs reduce BMP and ActA signaling by inducing degradation of the NSC’s constituent components. **a**, HEK293 cells were transiently transfected with either myc-ACVR1 or myc-ACVR1.R206H, in combination with HA-ACVR2B, and treated with ActA, or left untreated. Membranes were isolated, normalized for protein concentration, and proteins were immunoprecipitated using anti-myc magnetic beads. HA-ACVR2B co-immunoprecipitated with myc-ACVR1 or myc-ACVR1.R206H. ACVR1 and ACVR2B were detected using Myc and HA antibodies, respectively. Exposure to ActA resulted in a reduction of myc-ACVR1 and HA-ACVR2B in the membrane of cells expressing myc-ACVR1. Conversely, in cells expressing myc-ACVR1.R206H, both myc-ACVR1.R206H and ACVR2B levels do not change appreciably during exposure to ActA. **b**, *Bmpr1a^−/-^*; *Bmpr1b^−/-^* mESCs were either left untreated or treated with 10nM ActA for 1 hour. After washing off unbound ActA, cells were exposed to varying concentrations of BMP7 for either 30 minutes or 2 hours. Immunoblot analysis demonstrates that ActA significantly decreases BMP7-induced Smad1/5 phosphorylation. **c**, Blocking the binding of ActA to ACVR1 increases ActA availability and Smad2/3 signaling. W20 cells were treated with either 300nM isotype control antibody or ACVR1 blocking antibody for 1 hour, followed by the addition of varying concentrations of ActA for another hour. Immunoblot analysis demonstrates that ActA-induced Smad2/3 phosphorylation is increased in the cells treated with ACVR1 antibody.

One of the predictions of these results is that high levels of ActA will render cells unresponsive to BMP signaling via ACVR1 even after ActA has been removed. To explore this prediction, we tested the effect of ActA pretreatment on BMP7-initiated signaling in HEK293 cells coexpressing myc-ACVR1 and HA-ACVR2B. We treated cells for 60 minutes with 10nM ActA, removed any excess by washing with PBS, then introduced BMP7 and measured Smad1/5 phosphorylation. ActA pretreatment resulted in a severe reduction of Smad1/5 phosphorylation in response to BMP7, 30 minutes after its addition. The cells recovered their ability to respond by 2 hours after exposure to ActA (**Extended Data Fig. 7a, b**), presumably via replenishment of degraded receptors by newly synthesized ones.

To make certain that these results are not an artifact of overexpression, we also performed a similar experiment in mESCs. We utilized mESCs that are homozygous-null for both *Bmpr1a* and *Bmpr1b* (*Bmpr1a^−/-^*; *Bmpr1b^−/-^*) to avoid a competing Smad1/5 signal from activation of these type I receptors and followed the same treatment protocol as for HEK293 cells. ActA pretreatment resulted in severe reduction of Smad1/5 phosphorylation in response to BMP7, mirroring the data obtained in HEK293 cells. In mESCs that had been exposed to ActA, 10-fold more BMP7 was needed to achieve an equivalent level of signaling as that observed in cells that had not been exposed to ActA (**Fig. 2b**). We extended these observations in W20 cells, which naturally express higher levels of ACVR1 than mESCs along with comparable levels of BMPR1A (**Extended Data Fig. 7c**) and again followed the same treatment protocol. Consistent with results thus far, pretreatment of W20 cells with 10nM Activin limited their responsiveness to 10 nM BMP7 (**Extended Data Fig. 7d, e**). This effect was less pronounced with 50 nM BMP7 presumably due to signaling via BMPR1A ^30, 31^. These results verify our initial demonstration that ActA acts as an inhibitor of BMP signaling mediated by ACVR1 ^11^ but provide new mechanistic insight, that it is not merely occupancy of ACVR1•IIR heterocomplexes by ActA but rather their induced lysosomal degradation that underlies inhibition of BMP signaling. Furthermore, these results indicate that this inhibition can be temporally sustained until lysosomally degraded receptors have been replenished.

### Inhibition of NSC formation enhances ActA signaling via ACVR1B

Another predicted outcome of the lysosomal degradation of the components of **ACVR1•ActA•IIR** NSCs is that ActA’s availability will also be reduced. Hence, we hypothesized that inhibition of ActA’s interaction with ACVR1 (and its incorporation into NSCs) would enhance ActA signaling via ACVR1B. To test this hypothesis, we treated W20 cells, with an ACVR1 antibody (REGN5168) that blocks the interaction of ACVR1 with its ligands ^4^. Treatment with REGN5168 enhanced Smad2/3 signaling by ActA (**Fig. 3c**), consistent with a mechanism where blocking ActA’s incorporation into NSCs with ACVR1 increases its availability to interact with other proteins, such as ACVR1B.

**Figure 3.**
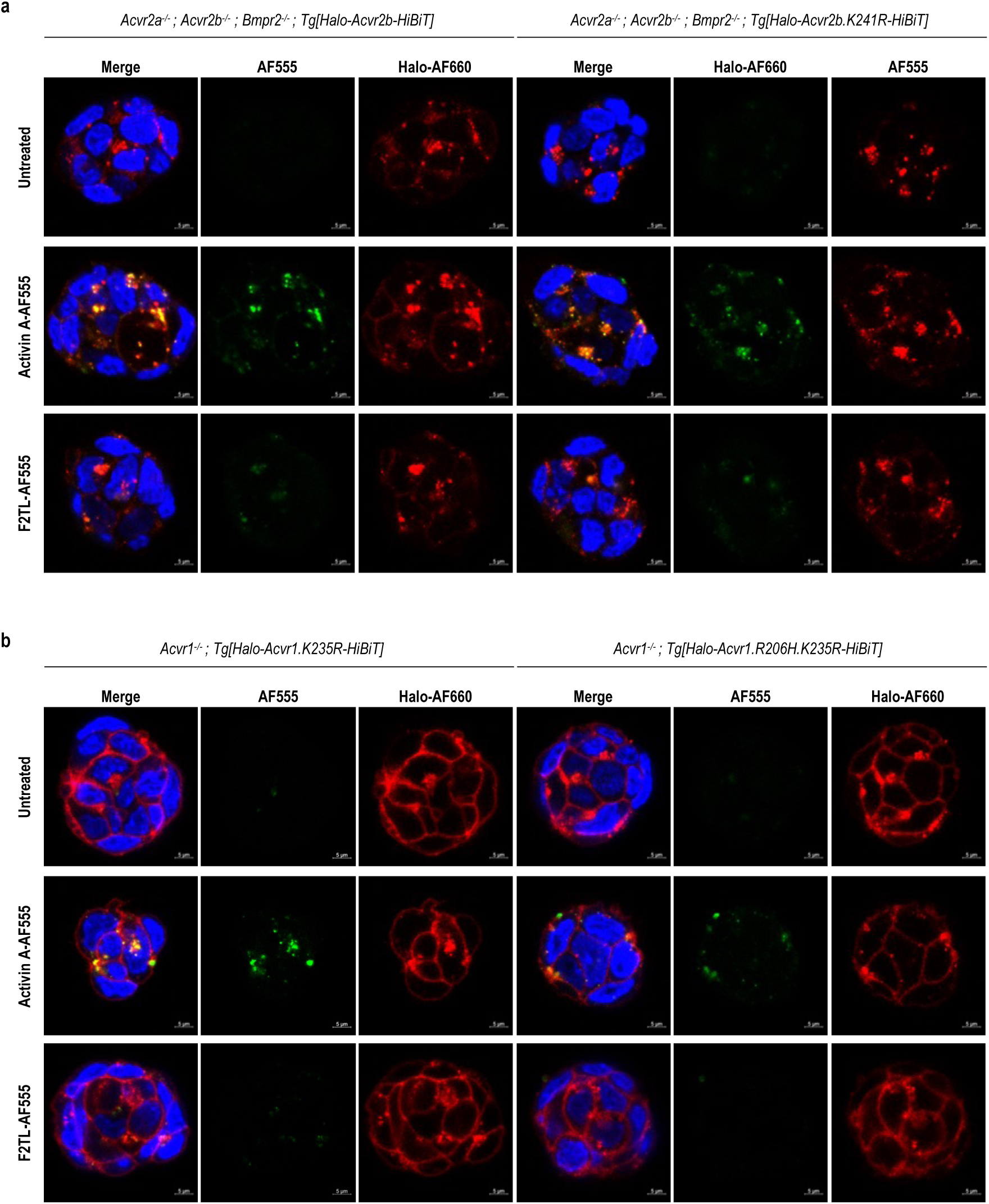
Signaling is not a determinant of NSC endocytosis. **a**,**b**, *Acvr2a^−/-^; Acvr2b^−/-^; Bmpr2^−/-^* mESCs expressing Halo-ACVR2B-HiBiT and Halo-ACVR2B.K241R-HiBiT (**a**), as well as *Acvr1^−/-^* mESCs expressing Halo-ACVR1.K235R-HiBiT and Halo-ACVR1.R206H.K235R-HiBiT (**b**) were either treated with 30nM ActA-AF555 or F2TL-AF555 (green) or left untreated. ACVR1 and ACVR2B proteins were visualized with AF660 Halo ligand (red) and DAPI (blue). Confocal microscopy was used for imaging. ActA-AF555 internalized in kinase-dead Halo-ACVR1.K235R-HiBiT, Halo-ACVR2B-HiBiT, and kinase-dead Halo-ACVR2B.K241R-HiBiT expressing mESCs, but not in kinase dead Halo-ACVR1.R206H.K235R-HiBiT expressing mESCs. F2TL-AF555 was not internalized in any of the tested mESC lines.

### NSC endocytosis requires interaction between ACVR1 and ActA

To understand the requirements for NSC endocytosis we first explored the role of ACVR1. We employed F2TL as it cannot engage ACVR1 directly, yet can drive dimerization of preformed (ligand-independent) heterodimers of ACVR1 with type II receptors ^4, 20^. When F2TL-AF555 is utilized as the ligand, Halo-ACVR1-HiBiT and F2TL fail to internalize (**Fig. 1a; Extended Data Fig. 5a**), indicating that direct interaction between ActA and ACVR1 is required for endocytosis of the resulting complex.

Next, we interrogated whether ACVR1B participates in the endocytosis of ActA. We generated mESCs that either lack ACVR1 and retain ACVR1B (*Acvr1^−/-^; Acvr1b^+/+^*) or retain ACVR1 and lack ACVR1B (*Acvr1^+/+^; Acvr1b^−/-^*) and followed endocytosis of ActA-AF555. Loss of ACVR1B does not impact endocytosis of ActA-AF555, and may even increase it, whereas loss of ACVR1 greatly reduces it (**Extended Data Fig. 8)**. These results indicate that ACVR1B is not required for NSC endocytosis and does not appear to participate in this process.

### NSC endocytosis requires type II receptors

To explore further the molecular mechanism by which NSCs carry out their function we asked whether type II receptors are required. Although ACVR1 is associated with type II receptors in non-covalent, ligand-independent heterodimeric complexes ^4, 32^, we asked whether the association of ACVR1 with ActA alone might be adequate to cause trafficking of the resulting complex to lysosomes, even though that association would be very transient ^20^. To determine this, we engineered mESCs that lack all three type II receptors that ActA has been shown to associate with, namely ACVR2A, ACVR2B, and BMPR2 (*Acvr2a^−/-^; Acvr2b^−/-^; Bmpr2^−/-^*), and tested internalization of ActA-AF555. Absence of these receptors result in a complete loss of internalization of ActA (**Extended Data Fig. 9**). When the expression of ACVR2B was restored via lentiviral transfer of Halo-ACVR2B-HiBiT to derive *Acvr2a^−/-^; Acvr2b^−/-^; Bmpr2^−/-^; Tg[Halo-Acvr2b-HiBiT]* mESCs (**Extended Data Fig. 1a, c, 3b, 10**), the response to ActA was restored, with both ActA and Halo-ACVR2B-HiBiT trafficking to the lysosome (**Fig. 3a, left panel**).

### Signaling is not a determinant of NSC internalization

Given that stoichiometry of the **ACVR1•ligand•IIR** complexes is not a determinant of endocytosis, we explored whether signaling might be that determinant. To eliminate variables that could be introduced by varying the ligand, we engineered a mESC line wherein ACVR1.R206H was rendered kinase dead (ACVR1.R206H.K235R). We hypothesized that if it is signaling that stops **ACVR1.R206H•ActA•IIR** complexes from trafficking to the lysosome, then disabling that signaling by rendering ACVR1.R206H kinase dead the resulting **ACVR1.R206H.K245R•ActA•IIR** complexes should mimic the NSCs formed with wild type ACVR1. Hence, we examined trafficking of Halo-ACVR1.R206H.K235R-HiBiT in *Acvr1^−/-^; Tg[Halo-Acvr1.R206H.K235R-HiBiT]* mESCs and compared it with its non-R206H kinase-dead counterpart, Halo-ACVR1.K235R-HiBiT in *Acvr1^−/-^; Tg[Halo-Acvr1.K235R-HiBiT]* mESCs. In the presence of ActA, Halo-ACVR1.K235R-HiBiT trafficked to lysosomes, whereas Halo-ACVR1.R206H.K235R-HiBiT remained on the surface (**Fig. 3b**). Hence, the complex of **ACVR1.R206H.K235R•ActA•IIR,** which is unable to signal, retains the properties of its signaling counterpart – **ACVR1.R206H•ActA•IIR** – rather than adopt the properties of **ACVR1•ActA•IIR** NSCs.

Similar results were obtained via pharmacological inhibition of ACVR1 kinase activity using the small molecule LDN214117 ^33^, which – when used at a concentration of 1 nM – very efficiently blocks ActA-induced Smad1/5/8 signaling from ACVR1.R206H while leaving activation of Smad2/3 signaling via ACVR1B unaffected (**Fig. 4a)**. Inhibition of ACVR1 kinase in *Acvr1^−/-^ ; Tg[Halo-Acvr1.R206H-HiBiT]* mESCs does not drive endocytosis of **ACVR1.R206H•ActA•IIR** complexes. As would be expected, LDN214117 treatment of *Acvr1^−/-^ ; Tg[Halo-Acvr1-HiBiT]* mESCs, does not affect lysosomal trafficking of **ACVR1•ActA•IIR** complexes (**Fig. 4b**).

**Figure 4.**
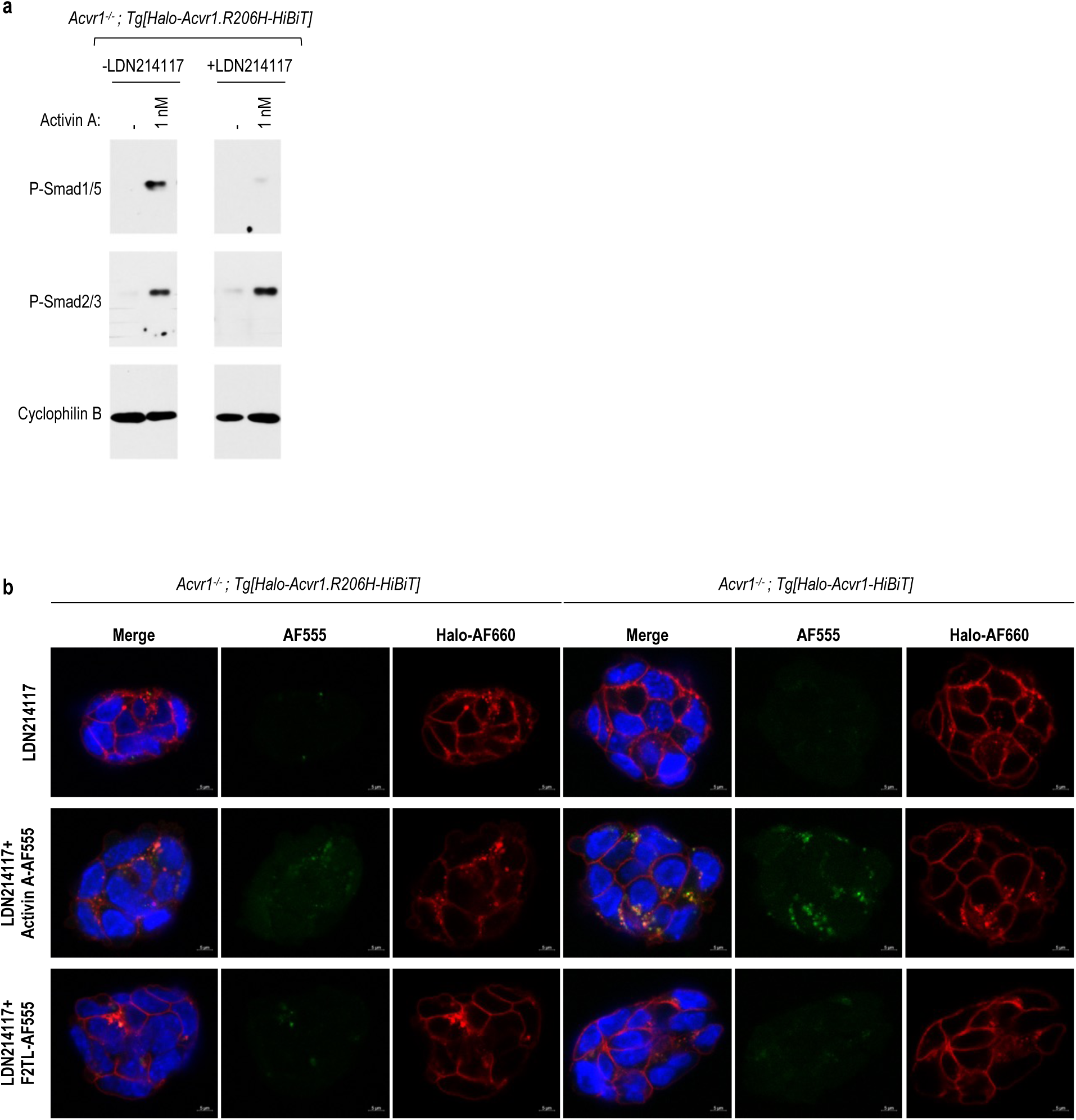
Inhibition of ACVR1.R206H kinase activity does not result in lysosomal trafficking of ACVR1.R206H•ActA•IIR complexes. **a**, *Acvr1^−/-^* mESCs expressing Halo-ACVR1.R206H-HiBiT were treated with 10nM LDN214117 for 1 hour followed by the addition of 1nM ActA for another hour. LDN214117 inhibited ActA-induced Smad1/5 phosphorylation, but not Smad2/3 phosphorylation. **b**, *Acvr1^−/-^* mESCs expressing Halo-ACVR1-HiBiT and Halo-ACVR1.R206H-HiBiT were either treated with 10nM LDN214117 for 1 hour, followed by the addition of 30nM ActA-AF555 or F2TL-AF555 (green). ACVR1 and ACVR2B proteins were visualized with AF660 Halo ligand (red) and DAPI (blue). Confocal microscopy was used for imaging. ActA-AF555 did not internalize in the LDN214117 treated Halo-ACVR1.R206H-HiBiT expressing mESCs but did internalize in the LDN214117 treated Halo-ACVR1-HiBiT mESCs. F2TL-AF555 was not internalized under either condition.

We also explored whether the kinase activity of type II receptors is required for endocytosis of NSCs. To this effect, we introduced Halo-ACVR2B-HiBiT or its kinase-dead counterpart, Halo-ACVR2B.K241R-HiBiT into *Acvr2a^−/-^; Acvr2b^−/-^; Bmpr2^−/-^* mESCs. The resulting lines were tested for responsiveness to BMP7. *Acvr2a^−/-^; Acvr2b^−/-^; Bmpr2^−/-^; Tg[Halo-ACVR2B-HiBiT]* mESCs respond to BMP7 whereas *Acvr2a^−/-^; Acvr2b^−/-^ ; Bmpr2^−/-^; Tg[Halo-ACVR2B.K241R-HiBiT]* mESCs do not **(Extended Data Fig. 10)**. We then tested their ability to internalize ActA-AF555. Lack of ACVR2B kinase activity had no obvious effect on the internalization of **ACVR1•ActA•IIR** NSCs (**Fig. 3a, right panel**). This data indicates that signaling is not a determinant of trafficking of NSCs to the lysosome; rather a different mechanism must be involved.

### The extracellular domain of ACVR1 is required for lysosomal trafficking but not for signaling

Given the tenuous interaction of ActA with ACVR1, plus the fact that F2TL (which does not bind ACVR1) can still drive the formation of signaling complexes of ACVR1.R206H with type II receptors ^20^, propelled us to test whether the extracellular domain of ACVR1 is required for either the formation of either **ACVR1•ActA•IIR** NSCs or **ACVR1•BMP7•IIR** signaling complexes. To do this, we introduced myc-tagged versions of ACVR1 or ACVR1.R206H and counterparts lacking the extracellular domain (Δecto) into cells lacking ACVR1 (*ACVR1^−/-^*) and tested their response to BMP7 and ActA. Both ACVR1Δecto and ACVR1Δecto.R206H retained the signaling properties of their full-length counterparts, with ACVR1Δecto responding to BMP7 and ACVR1Δecto.R206H responding to both BMP7 and ActA, though perhaps not as efficiently (**Fig. 5a, b**). This indicated that the extracellular domain of ACVR1 is dispensable for signaling. The kinase dead versions of the introduced full-length ACVR1 and ACVR1.R206H did not respond to either ligand, indicating that the signal observed is solely attributable to the introduced ACVR1 constructs.

**Figure 5.**
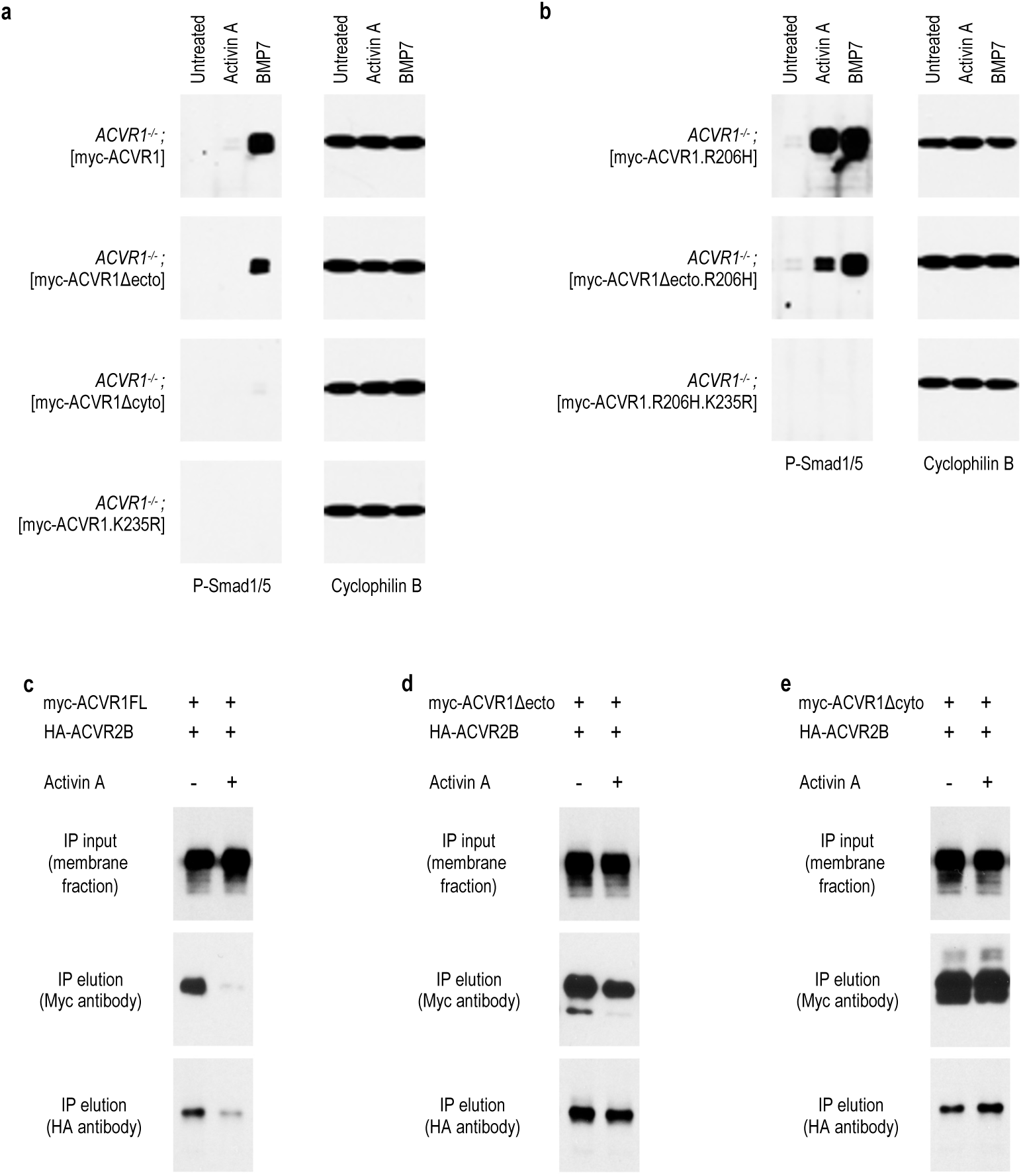
The extracellular domain of ACVR1 is dispensable for signaling but is required for induction of lysosomal trafficking. Different versions of myc-ACVR1 or myc-ACVR1.R206H (full length, Δecto, and K235R) as well as myc-ACVR1Δcyto were expressed in HEK293 cells that are null for endogenous ACVR1 (*ACVR1^−/-^*). The cells were treated with either ActA or BMP7 and activation of Smad1/5/8 signaling was visualized by Western blotting to pSmad1/5. **a**, BMP7 activates signaling from full length ACVR1 as well as ACVR1Δecto, whereas ActA fails to activate signaling. **b**, Both ActA and BMP7 activate signaling of ACVR1.R206H and ACVR1.R206HΔecto. The kinase-dead versions of ACVR1 and ACVR1.R206H, mycACVR1.K235R and ACVR1.R206H.K235R cannot activate signaling (**a**, **b**). **c**, **d**, **e**, HEK293 cells expressing HA-ACVR2B together with full length myc-ACVR1 (myc-ACVR1FL) or myc-ACVR1Δecto, or mycACVR1Δcyto were treated with ActA and the fate of these receptors was visualized. Whereas myc-ACVR1FL along with HA-ACVR2B are degraded when exposed to ActA (**c**), no such effect is observed for myc-ACVR1Δecto (**d**) and myc-ACVR1Δcyto (**e**).

We next examined whether lack of the extracellular domain affected lysosomal trafficking. For this, we utilized HEK293 cells coexpressing myc-tagged ACVR1 (myc-ACVR1), or ACVR1Δecto (myc-ACVR1Δecto), along with HA-tagged ACVR2B (HA-ACVR2B). Additionally, we tested whether the cytoplasmic domain of ACVR1 is required for lysosomal trafficking by introducing ACVR1 lacking its cytoplasmic (myc-ACVR1Δcyto). Consistent with the results shown above, treatment with ActA drives degradation of full-length ACVR1 along with ACVR2B (**Fig. 5c**). Conversely, in cell expressing ACVR1Δecto or ACVR1Δcyto, neither one of these two proteins nor ACVR2B is degraded (**Fig. 5d, e**). These results indicate that both the extracellular and the cytoplasmic domain of ACVR1 are required for trafficking to lysosomes.

### Other Activins also form NSCs with ACVR1 and suppress BMP signaling

Since Activin AB, B, and AC can activate signaling of ACVR1.R206H ^11, 12^, we surmised that those ligands must also form NSCs with wild type ACVR1. Hence, we tested whether Activin AB, B, and AC inhibit BMP7-induced signaling via ACVR1, using HEK293 cells stably expressing ACVR1. Consistent with prior observations ^11, 12^, we verified that Activin AB, B, and AC activate ACVR1.R206H (**Fig. 6a**). Having thus qualified that these ligands interact with ACVR1, we tested whether they could inhibit BMP7-induced signaling mediated by wild type ACVR1. Activin AB and B repressed BMP7-induced signaling similarly to Activin A, whereas Activin AC was less efficient (**Fig. 6b**). Next, we tested whether ACVR1•ligand•IIR NSCs formed with these Activins also trafficked to lysosomes. As described above, we utilized HEK293 cells stably expressing myc-ACVR1 and HA-ACVR2B, to enable evaluation of these receptors by Western blotting. Whereas exposure of these cells to ActA results in near complete degradation of myc-ACVR1 and HA-ACVR2B (**Fig. 2a**), such effect was not observed with the other Activins (**Fig. 6c**). Hence, although these other Activins form NSCs with ACVR1, these NSCs do not appear to induce internalization of ACVR1 at an appreciable level. This data implied that ActA is likely to have more pronounced and sustained effect on inhibiting BMP7 signaling through ACVR1 compared to other Activins. To test this, we utilized *Bmpr1a^−/-^*; *Bmpr1b^−/-^* mESCs, where ACVR1 is not overexpressed, and where activation of Smad1/5/8 signaling through BMPR1A and BMPR1B is avoided. As in the experiments described above, mESCs were treated with the different Activins for 60 minutes, then excess ligands were removed with washing, and subsequently the cells were exposed to BMP7 for 30 minutes. Although all the Activins suppressed BMP7-induced signaling, ActA was more efficient (**Fig. 6d**). This result is consistent with the notion that ActA causes sustained inhibition of BMP signaling mediated by ACVR1 because it causes lysosomal degradation of ACVR1 along with accompanying type II receptors. Furthermore, these results indicate that although ACVR1 can form NSCs with several other Activins, these NSCs differ qualitatively from to those formed with ActA as they do not result in lysosomal trafficking and degradation of the corresponding complexes.

**Figure 6.**
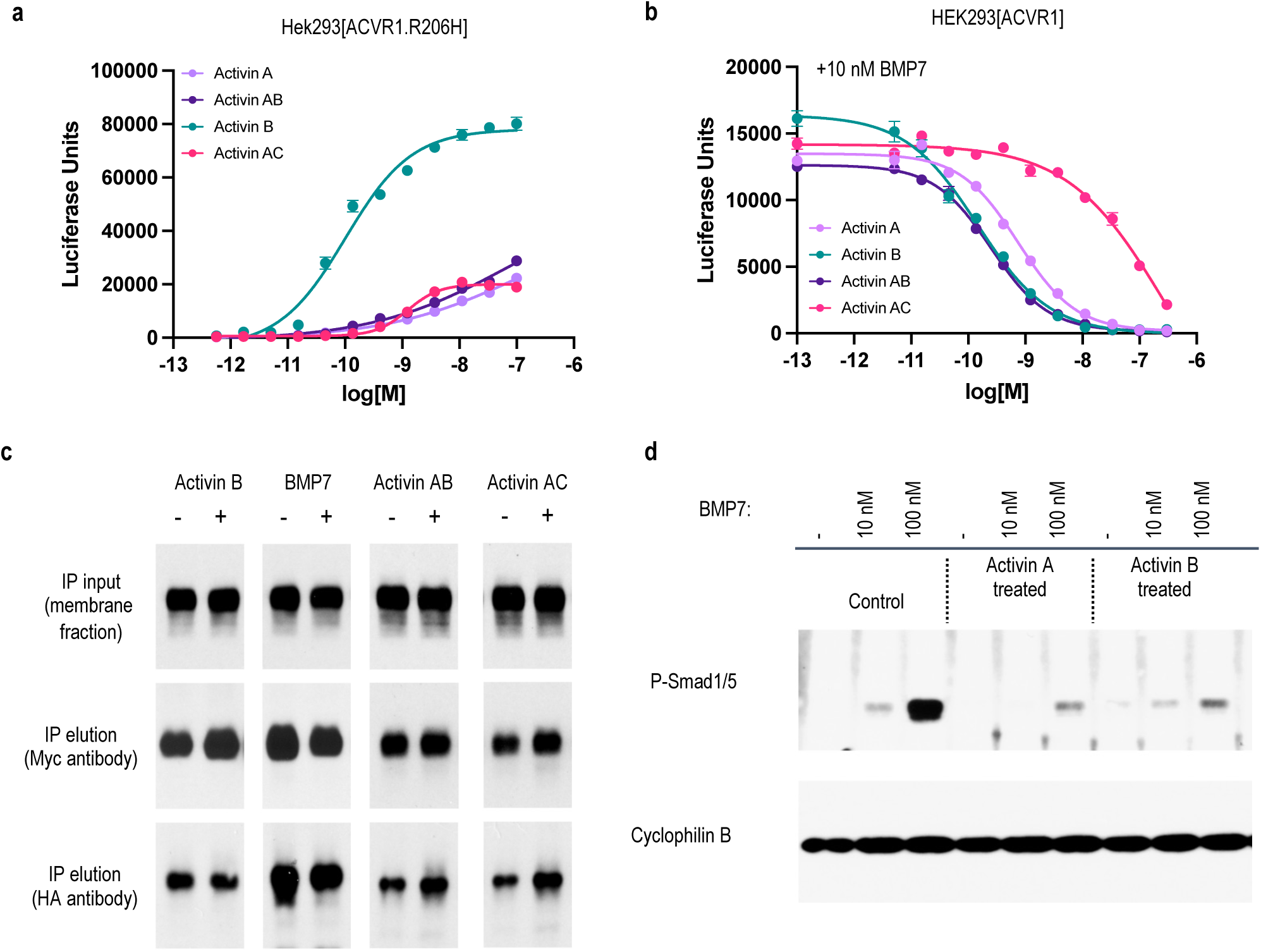
Other Activins also form NSCs with ACVR1 and suppress BMP signaling but do not drive lysosomal trafficking. **a**, ActA, Activin B, Activin AB, and Activin AC were titrated in a HEK293-BRE luciferase reporter cells (Smad1/5 responsive) overexpressing ACVR1.R206H. All these Activins induced Smad1/5 signaling in cells expressing the FOP mutant receptor ACVR1.R206H. **b**, HEK293-BRE luciferase reporter cells overexpressing wild type ACVR1 were treated with a fixed (10nM) concentration of BMP7, and different concentrations of ActA, Activin B, Activin AB, or Activin AC. Activins A, B, AB, and AC inhibited BMP7-induced Smad1/5 signaling to varying degrees, with Activin AC being the least efficient. **c**, HEK293 cells co-expressing myc-ACVR1 and HA-ACVR2B were treated with Activin B, BMP7, Activin AB, or Activin AC and the fate of these receptors was visualized. None induced degradation of myc-ACVR1 and HA-ACVR2B, indicating that although Activin B, Activin AB, or Activin AC are form NSCs with ACVR1 and ACVR2B, these complexes do not traffic to lysosomes at an appreciable rate. **d**, *Bmpr1a^−/-^*; *Bmpr1b^−/-^* mESCs were either left untreated or treated with 10nM of ActA or Activin B for 1 hour. After washing off any unbound Activins, these cells were exposed to varying concentrations of BMP7 for 30 minutes. Both ActA and Activin B blocked BMP7-induced Smad1/5 phosphorylation, but the effect of ActA was more pronounced.

## Discussion

We report here a novel regulatory mechanism in the BMP/TGFß pathway. ActA and ACVR1 coordinately regulate their own signaling by forming **ACVR1•ActA•IIR** NSCs that rapidly traffic to lysosomes (**Fig. 7**. Conversely, identical complexes utilizing the FOP-causing variant ACVR1.R206H remain on the membrane and signal. This stands in contrast to another Activin receptor, ACVR1B, which appears to cycle between the membrane and early endosomes irrespective of engagement by ligand ^34^.

**Figure 7.**
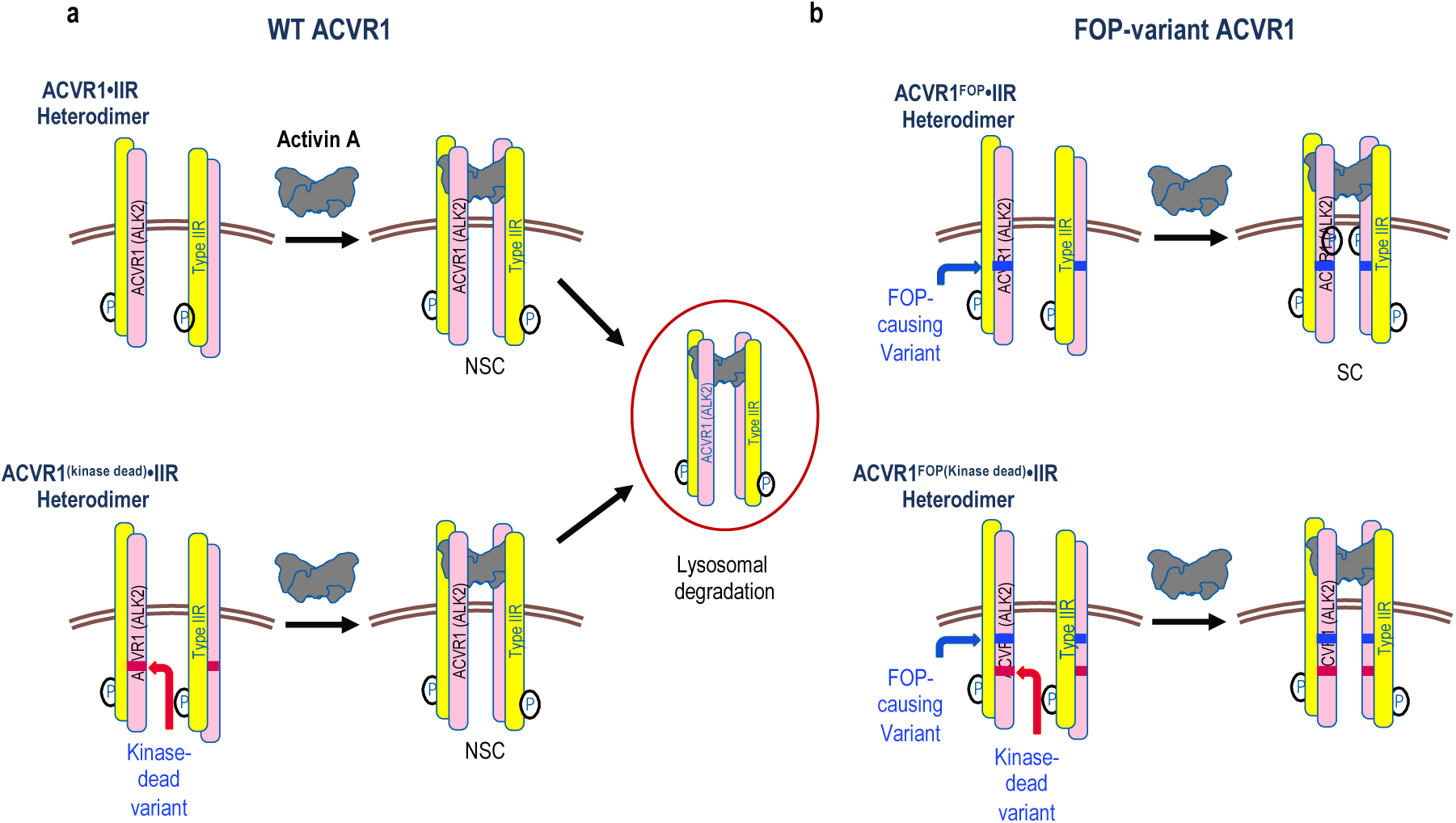
Activin and ACVR1 coordinately regulate their own availability. **a**, When ActA binds wild type ACVR1 it drives the formation of NSCs that traffic to lysosomes wherein the components of the NSC are degraded. This results in a reduction of the NSC’s constituent components and hence their availability for signaling. Trafficking of NSCs to the lysosome does not require kinase activity of either ACVR1 or IIR (not depicted). **b**, When ActA engages FOP-variant ACVR1 such as ACVR1.R206H, the resulting complexes remain on the membrane and signal (SC). Inhibitin signaling of ACVR1.FOP•ActA•IIR complexes for example by rendering ACVR1.FOP kinase dead does not enable trafficking of such complexes to lysosomes, indicating that signaling is not a determinant of this process.

In the lysosome, the components of **ACVR1•ActA•IIR** NSCs are degraded, resulting in a reduction in ACVR1 and associated type II receptors ^4^ on the cell surface. Consequently, in the presence of ActA, BMP signaling through ACVR1 is reduced. This effect can be temporally sustained, with cells treated with ActA remaining unresponsive to subsequent stimulation by BMPs even after ActA has been removed, and presumably until ACVR1 and type II receptors have been replenished by de novo biosynthesis. The degree to which BMP signaling is suppressed depends on the expression level of ACVR1 relative to other type I BMP receptors: in cells where the main type I BMP receptor is ACVR1, signaling by BMPs is greatly inhibited, whereas in cells where other type I BMP receptors are expressed BMP signaling is less subject to the effects of NSCs.

Furthermore, NSCs appear to be a mechanism that limits the amount of available ActA. In cells where ACVR1 is highly expressed in comparison to ACVR1B, the responsiveness of the latter to ActA is reduced. Hence, inhibition of binding of ActA to ACVR1 results in higher levels of signaling via ACVR1B. As antagonism is reciprocal, NSCs have the capacity to most potently regulate the more limiting component – ACVR1•IIR heterodimers and BMP signaling versus ActA and its activation of ACVR1B. Our observations agree with a mathematical model showing that the outcome of signaling by any given set of ligands depends on the specific repertoire of receptors (both with respect to their identity and their relative abundance) in the responding cell ^35^. Similar observations pertain to Activin AB, B, and AC, as they also form NSCs with ACVR1, and they demonstrate that unlike most other BMP/TGFß family members which act solely as agonists or antagonists of signaling, these Activins have a dual role as both agonists of Smad2/3 signaling through ACVR1B or TGFBR1 and antagonists of Smad1/5 signaling through ACVR1. However, unlike ActA, the NSCs that ACVR1 forms with Activin AB, B, and AC do not traffic to lysosomes. Perhaps this explains why these other Activins are not as efficient in mediating as temporally sustained a repression of BMP signaling as ActA.

From a molecular mechanism viewpoint, it is an open question how the complexes of ACVR1 with ActA drive lysosomal trafficking whereas the complexes with the other Activins do not. Likewise, the broader question of how stoichiometrically identical ACVR1•ligand•IIR complexes bring about such different outcomes, defined by the identity of the ligand remains unanswered. The finding that the extracellular domain of ACVR1 is required for lysosomal trafficking of the ACVR1 NSCs with ActA (but is dispensable for signaling with BMPs) indicates that very specific interactions between different Activins and ACVR1 govern the properties of the corresponding NSCs and suggests ligand-specific conformational changes as a potential mechanism.

Our findings have implications for therapeutic approaches targeting ACVR1 signaling in FOP and other indications ^36–38^. Approaches under investigation ^39^ include inhibition of ActA ^15^ or ACVR1 ^40^ using blocking antibodies, inhibition of ACVR1 kinase ^36, 41–43^, and siRNA-mediated reduction of ACVR1 expression ^44^. Our findings predict that at the molecular level these approaches will have different outcomes. For example, blocking ActA with an antibody will not only abrogate activation of ACVR1.FOP, but also block the formation of **ACVR1•ActA•IIR** NSCs, whereas it will leave signaling by BMPs via ACVR1 unperturbed, and potentially enhance responsiveness. Inhibition of ACVR1’s kinase activity will block signaling via ACVR1 (whether FOP-variant or wild type) but will leave NSCs unperturbed; notably, inhibition of ACVR1.FOP will not convert **ACVR1.FOP•ActA•IIR** complexes into NSCs. Inhibition of ACVR1 using antibodies will inhibit the formation of signaling and non-signaling complexes alike; however, it is our assessment that ACVR1 antibodies should not be considered as therapeutics for FOP ^4, 16, 45, 46^. Lastly, methods that decrease the expression of ACVR1, whether it is via siRNA, ASO, or gene editing will impact all ACVR1’s functions equally unless they are allele specific.

These considerations are potentially important, given that the physiological roles of NSCs outside of FOP remain to be elucidated. Loss of *Acvr1* ^47–52^ or *Inhba* ^53^ result in multiple phenotypes. It is possible that a subset of these phenotypes may result from loss of NSC function rather than loss of signaling. We propose that elucidation of the molecular mechanisms driving phenotypes resulting from genetic alterations in either ACVR1 or any of the NSC-forming Activins should take into consideration both the ability of these proteins to acts as signaling molecules as well as their ability to suppress BMP signaling and/or regulate their own availability. Given that other BMP/TGFß family members have been described to act as antagonists of signaling ^5–8^, our results also call for a more careful exploration of the molecular behavior of the corresponding complexes.

## EXTENDED DATA

**Extended Data Figure 1.**
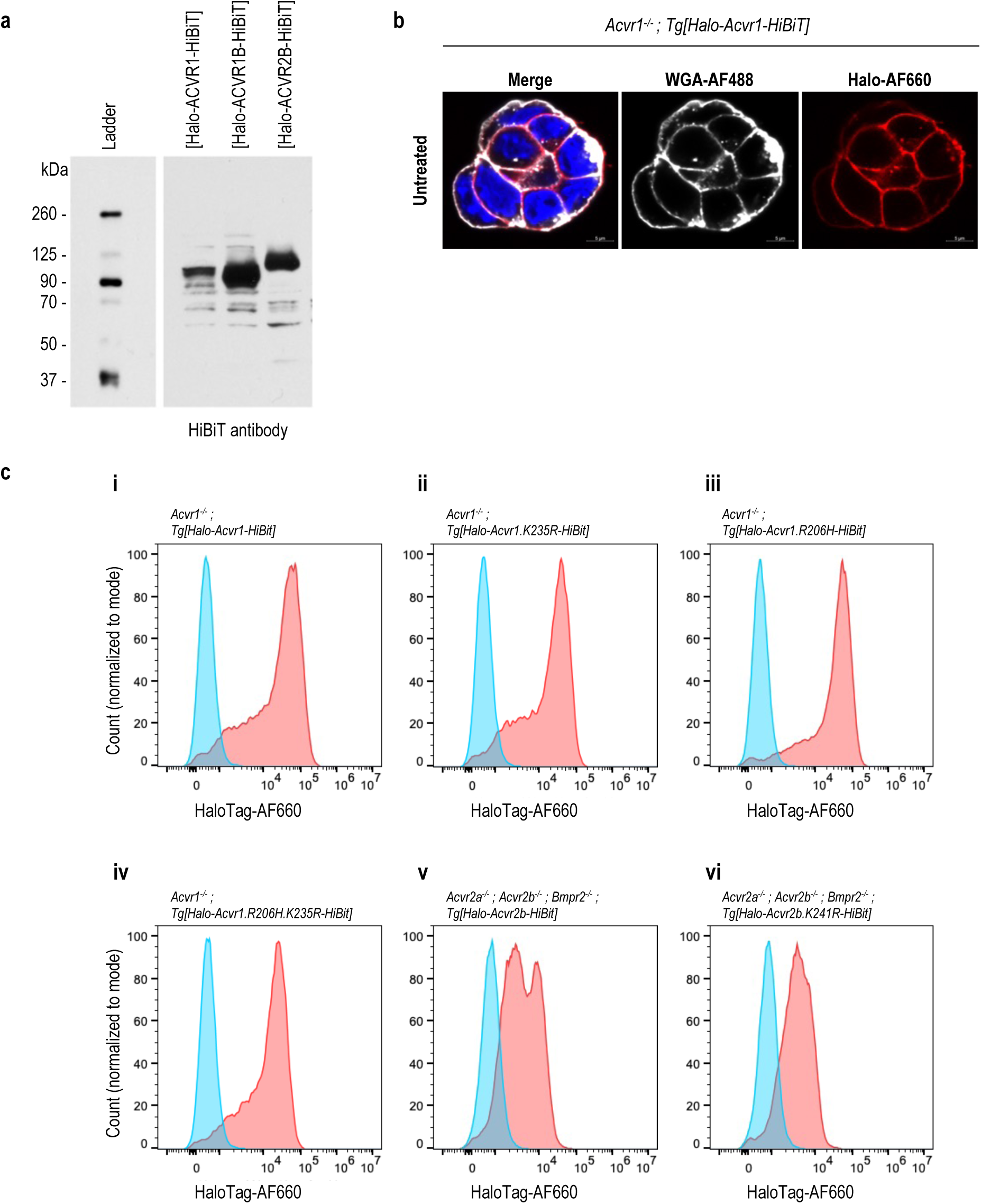
Expression of N-terminally Halo and C-terminally HiBiT tagged receptors in mESCs. **a**, Western blot of Halo-ACVR1B-HiBiT, Halo-ACVR1B-HiBiT, and Halo-ACVR2B-HiBiT. 10µg cell lysate from *Acvr1^−/-^* mESCs expressing Halo-ACVR1-HiBiT, *Acvr1b^−/-^* mESCs expressing Halo-ACVR1B-HiBiT and *Acvr2a^−/-^*; *Acvr2b^−/-^*; *Bmpr2^−/-^*mESCs expressing Halo-ACVR2B-HiBiT were resolved on an SDS-PAGE. The presence of these proteins was detected using a HiBiT antibody. **b**, Halo-ACVR1 localizes on the membrane. *Acvr1^−/-^* mESCs expressing Halo-ACVR1-HiBiT were stained with AF660 Halo ligand (red) and fluorescently labeled wheat germ agglutinin WGA-AF488 (white) to mark the Halo-ACVR1-HiBit receptor and cell membrane, respectively. Nuclei staining is with DAPI (blue). Confocal microscopy was used for imaging. Co-localization of the Halo tag and WGA staining demonstrates that Halo-ACVR1-HiBiT localizes on the cell membrane. **c**, Cell surface expression of Halo-ACVR1B-HiBiT and Halo-ACVR2B-HiBiT and variants thereof. *Acvr1^−/-^* mESCs expressing Halo-ACVR1-HiBiT **(i)**, Halo-ACVR1.K235R-HiBiT **(ii)**, Halo-ACVR1.R206H-HiBiT **(iii)**, Halo-ACVR1.R206H.K235R-HiBiT **(iv)**, *Acvr2a^−/-^*; *Acvr2b^−/-^*; *Bmpr2^−/-^*mESCs expressing Halo-ACVR2B-HiBiT **(v)**, and Halo-ACVR2B.K241R-HiBiT **(vi)** were stained with AF660 Halo ligand and analyzed by flow cytometry. All the tested cell lines showed positive receptor expression on the cell surface. Red curves represent AF660 Halo ligand-stained cells, and blue curves represent unstained cells.

**Extended Data Figure 2.**
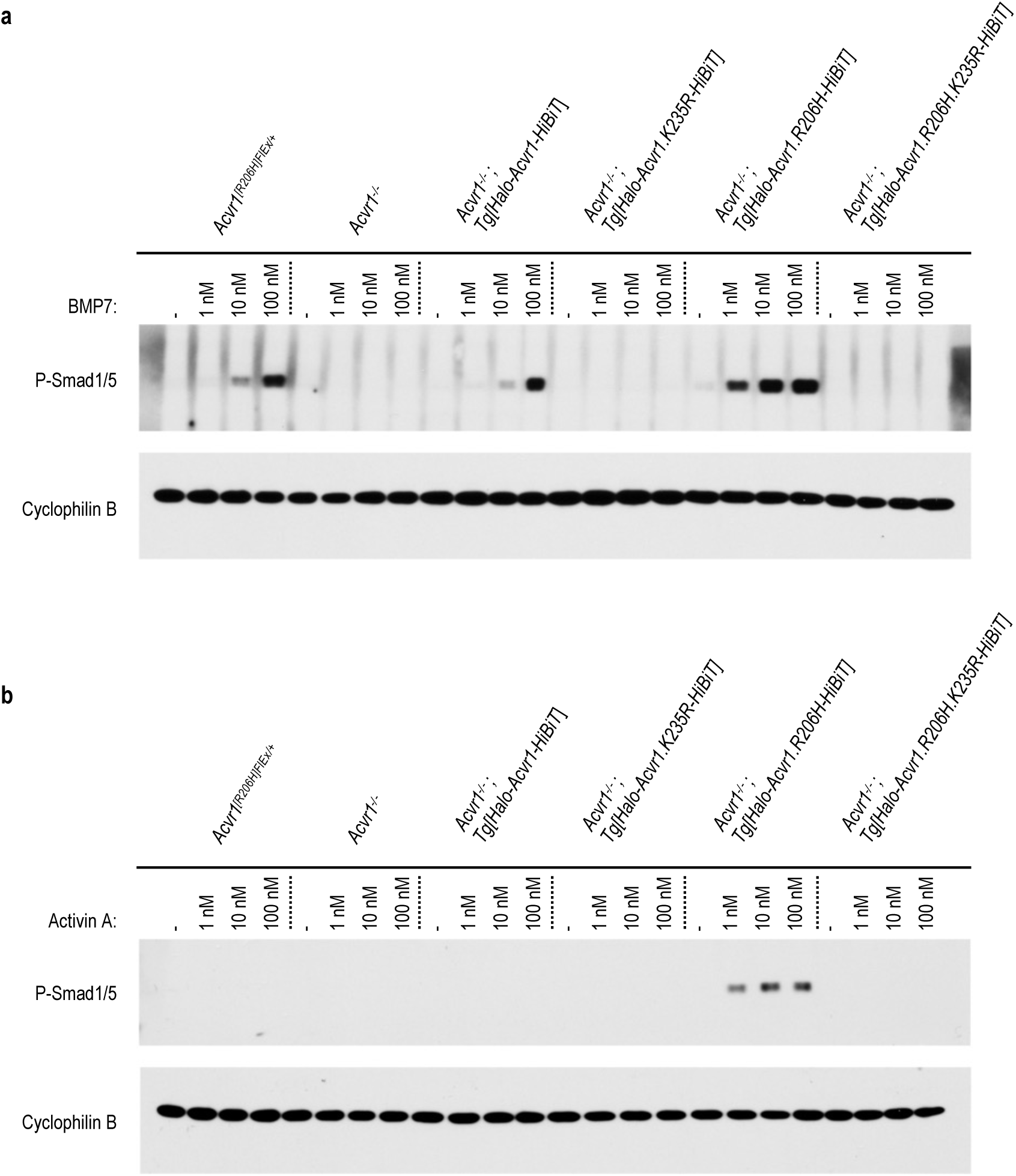
Ligand responsiveness of N-terminally Halo and C-terminally HiBiT tagged ACVR1 and variants thereof. **a**,**b**, *Acvr1^[R206H]FlEx/+^* mESCs, *Acvr1^−/-^* mESCs*, Acvr1^−/-^* mESCs expressing Halo-ACVR1-HiBiT, Halo-ACVR1.K235R-HiBiT, Halo-ACVR1.R206H-HiBiT, and Halo-ACVR1.R206H.K235R-HiBiT were treated with varying concentrations of BMP7 (**a**) or ActA (**b**) for 1 hour. BMP7 induced Smad1/5 phosphorylation in the parental *Acvr1^[R206H]FlEx/+^* mESCs, mESCs expressing either wild-type Halo-ACVR1-HiBiT and FOP mutant Halo-ACVR1.R206H-HiBiT, but not in their kinase-dead counterparts, Halo-ACVR1.K235R-HiBiT and ACVR1.R206H.K235R-HiBiT. ActA induced Smad1/5 phosphorylation only in the FOP mutant Halo-ACVR1.R206H-HiBiT expressing mESCs, but not in its kinase-dead counterpart.

**Extended Data Figure 3.**
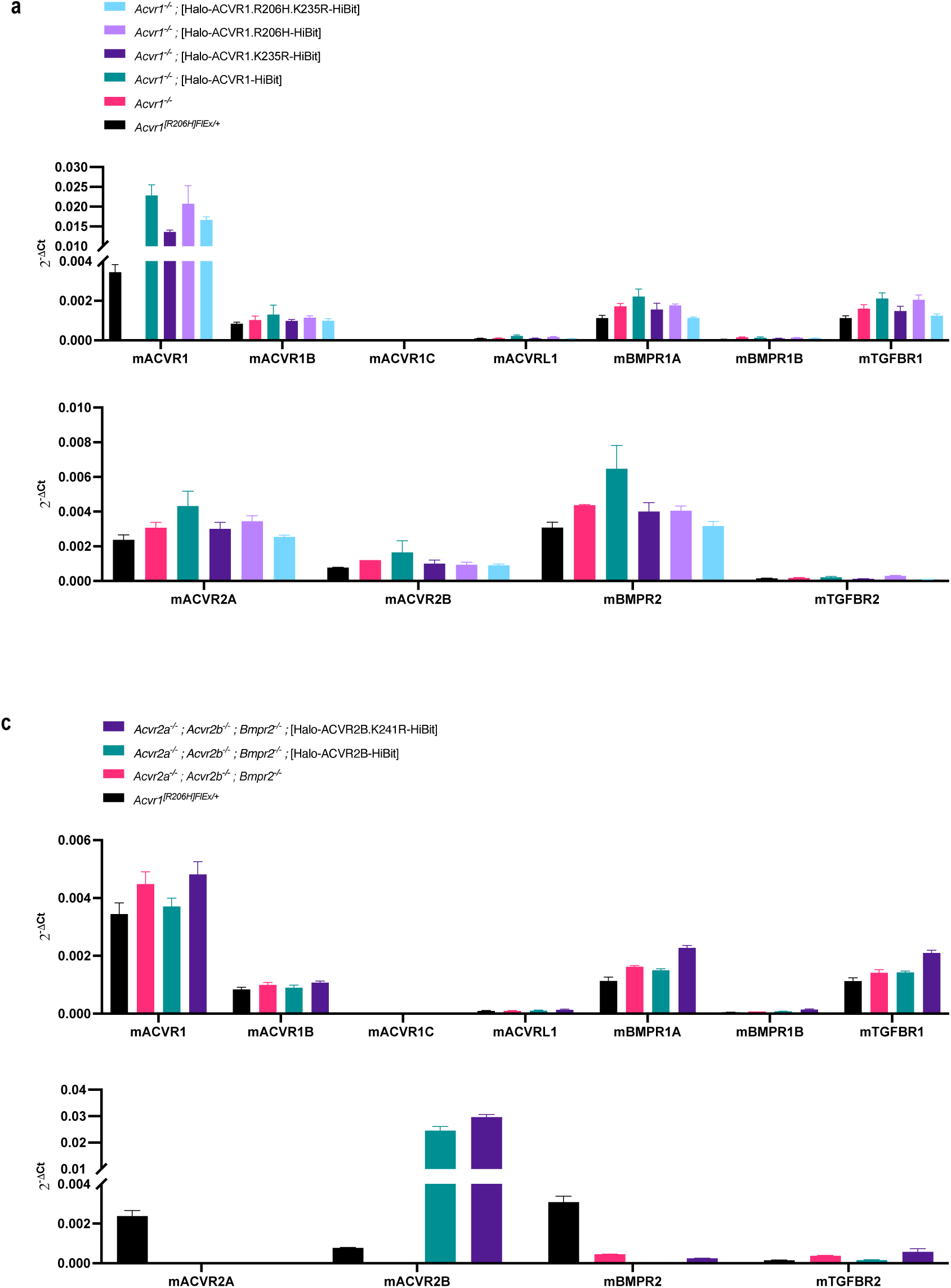
Expression levels of type I and type II BMP and TGFß receptors in Acvr1^−/-^ mESCs transgenically expressing Halo-ACVR1-HiBiT or Halo-ACVR2B-HiBiT and variants thereof. **a**, Type I and type II receptor mRNA levels of parental Acvr1[R206H]^FlEx/+^ mESCs, Acvr1^−/-^ mESCs, Acvr1^−/-^; Tg[Halo-ACVR1-HiBiT] mESCs , Acvr1^−/-^; Tg[Halo-ACVR1.K235R-HiBiT] mESCs, Acvr1^−/-^; Tg[Halo-ACVR1.R206H-HiBiT] mESCs, and Acvr1^−/-^; Tg[Halo-ACVR1.R206H.K235R-HiBiT] mESCs. **b**, Type I and type II receptor mRNA levels of Acvr2a^−/-^; Acvr2b^−/-^; Bmpr2^−/-^ mESCs, Acvr2a^−/-^; Acvr2b^−/-^; Bmpr2^−/-^; Tg[Halo-ACVR2B-HiBiT] mESCs, and Acvr2a^−/-^; Acvr2b^−/-^; Bmpr2^−/-^; Tg[Halo-ACVR2B.K241R-HiBiT] mESCs. All the N-terminally Halo and C-terminally HiBiT tagged receptors are expressed in the corresponding mESC lines. Expression of the tagged receptors does not affect the mRNA levels of other receptors of the BMP/TGFß family.

**Extended Data Figure 4.**
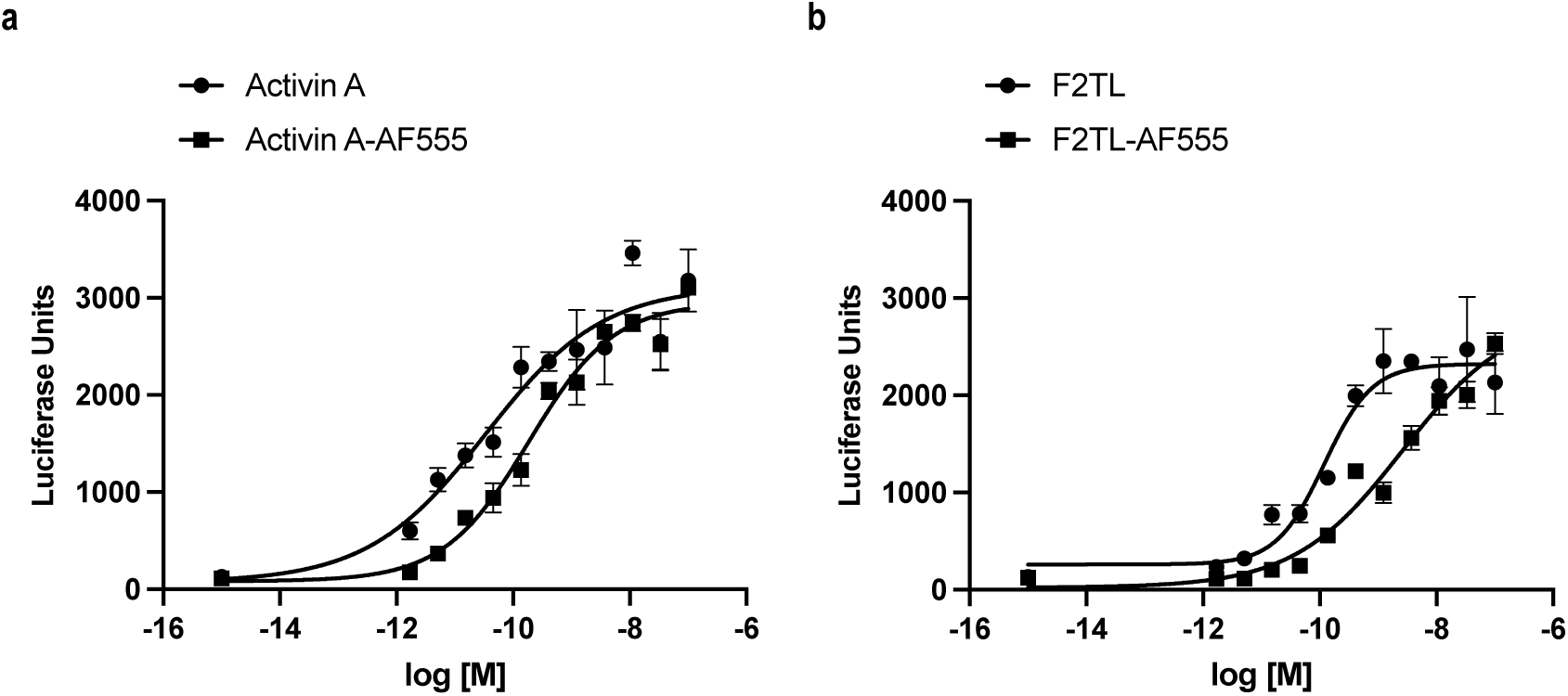
ActA-AF555 and F2TL-AF555 activate Smad2/3 signaling in HEK293-CAGA reporter cells. **a**,**b**, Activation of Smad2/3 signaling by both unlabeled and Alexa Fluor-555 (AF555) labeled ActA and F2TL was assessed in HEK293-CAGA luciferase reporter cells (responsive to Smad2/3 signaling). ActA, ActA-AF555 (**a**), and F2TL, F2TL-AF555 (**b**) were titrated in the HEK293-CAGA reporter cells. Both unlabeled and AF555 labeled ActiA and F2TL induce Smad2/3 signaling to a similar level.

**Extended data Figure 5.**
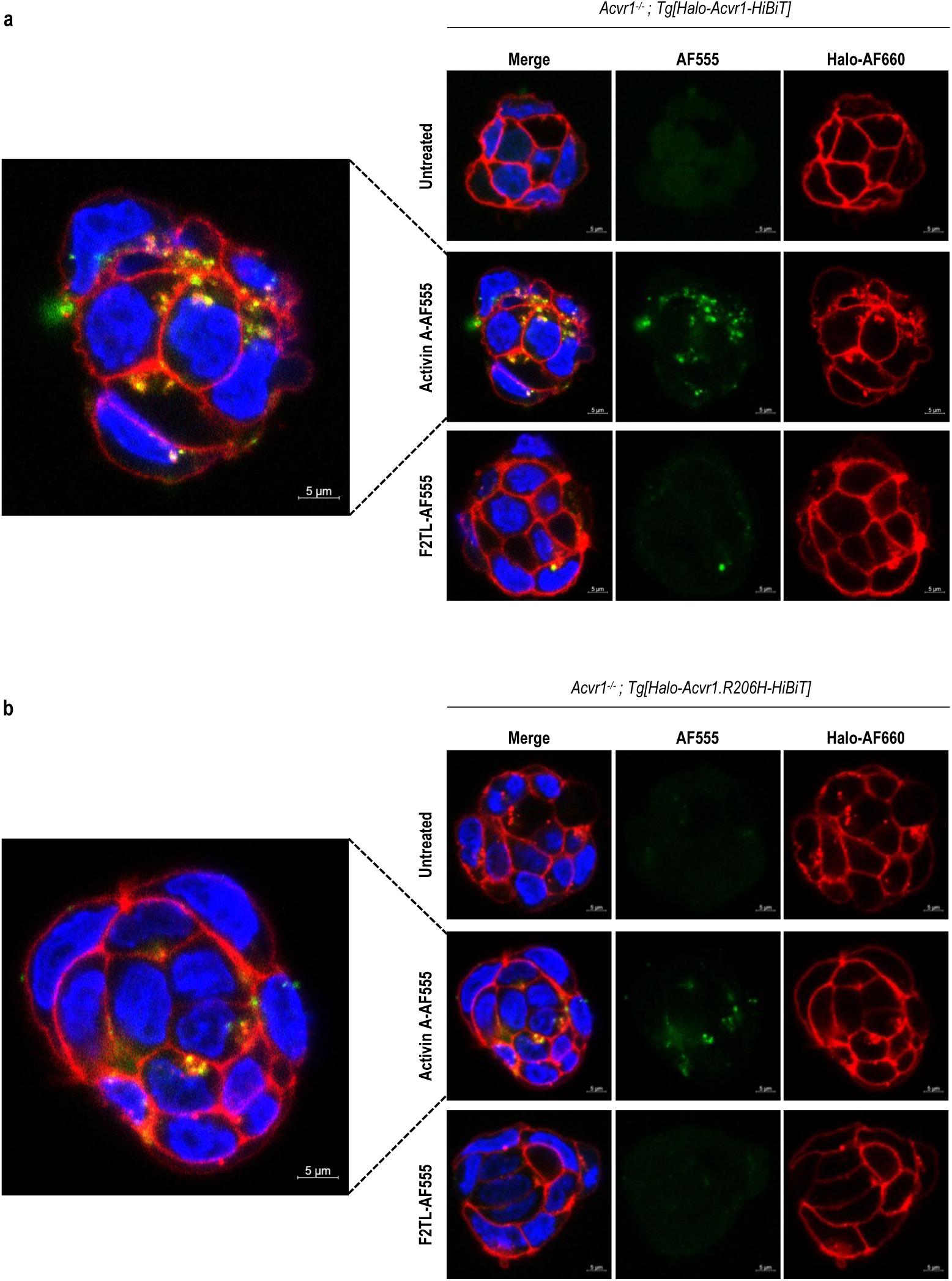
ACVR1•Activin A•IIR NSCs undergo endocytosis. **a**,**b**, Acvr1^−/-^ mESCs expressing either Halo-ACVR1-HiBiT (**a**) or Halo-ACVR1.R206H-HiBiT (**b**) were treated with 30nM ActA-AF555 or F2TL-AF555 or left untreated. ACVR1 proteins were visualized using AF660 Halo ligand (red). Nuclei were visualized using DAPI (blue). ActA or F2TL were detected via the AF555 fluorophore (green). Confocal microscopy was used for imaging. ActA-AF555 co-internalized with Halo-ACVR1-HiBiT but did not internalize with Halo-ACVR1.R206H-HiBiT. F2TL-AF555 did not drive endocytosis of either ACVR1 or ACVR1.R206H.

**Extended Data Figure 6.**
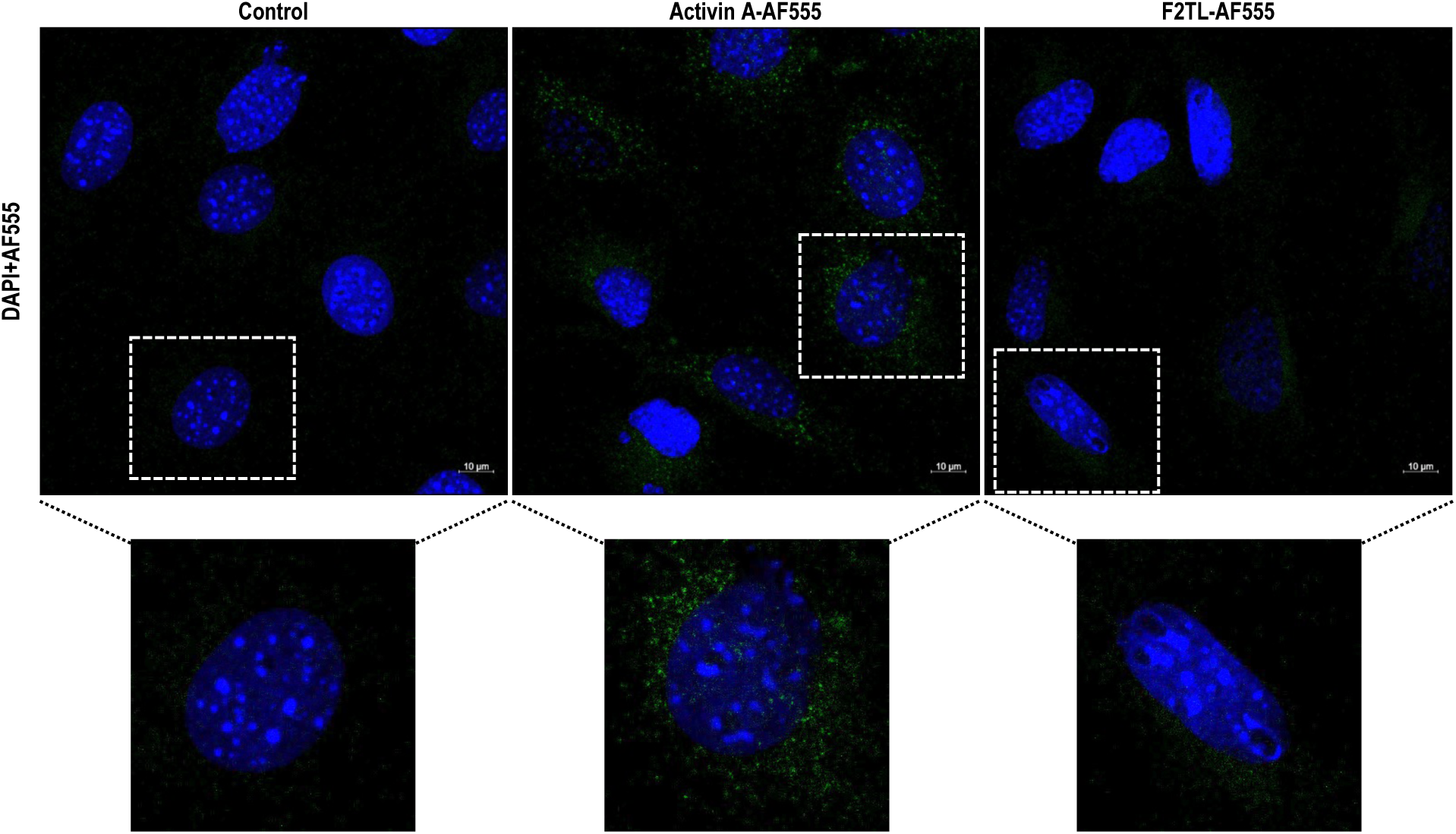
ActA, but not F2TL, undergoes endocytosis in W20 cells. W20 cells were treated with 30nM ActA-AF555 or F2TL-AF555 or left untreated. The cells were then stained with DAPI (blue) to visualize the cell nuclei. Confocal microscopy was used for imaging, and images from the Z-stack are shown. ActA-AF555 was endocytosed in W20 cells, but F2TL-AF555 did not.

**Extended Data Figure 7.**
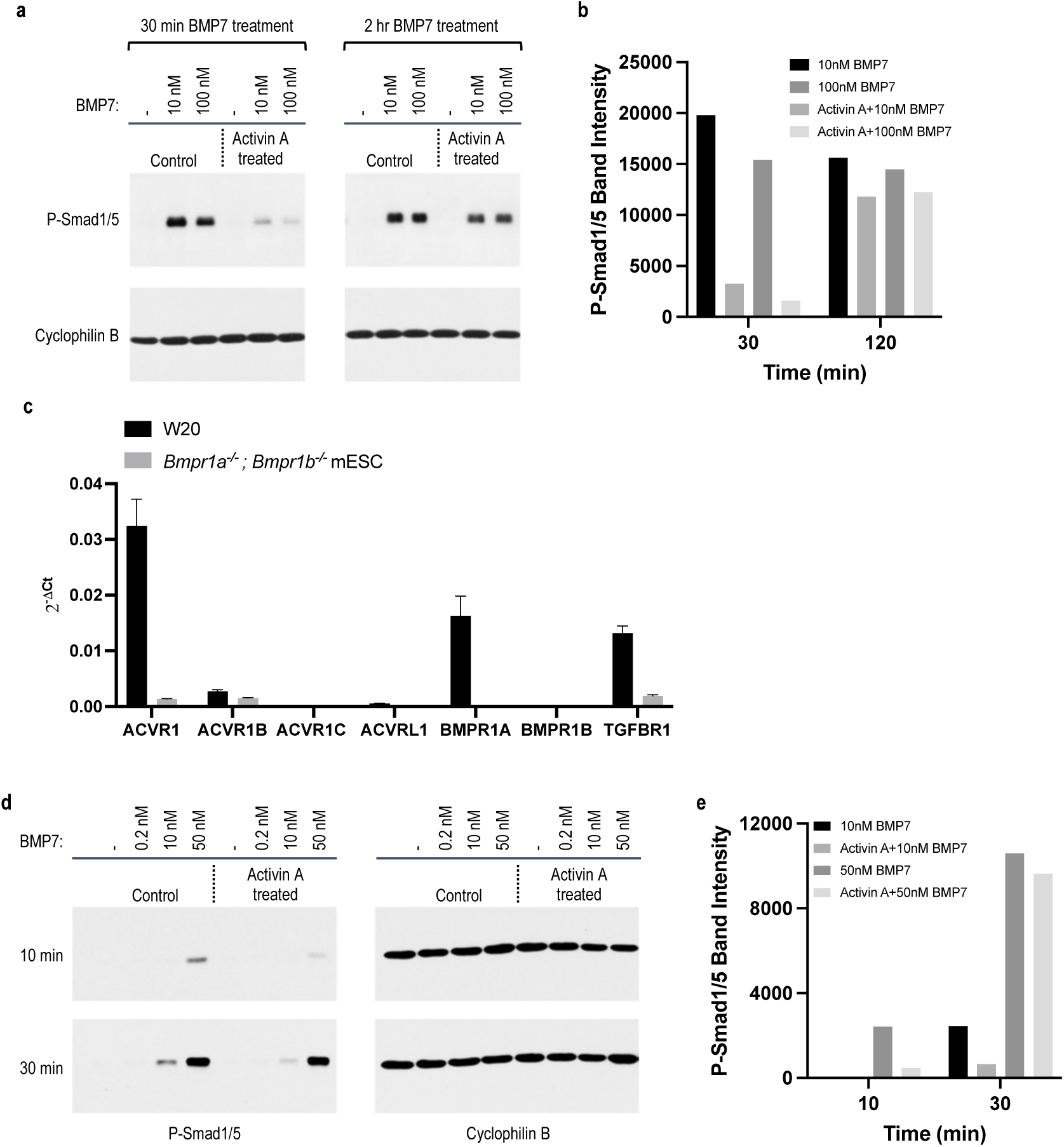
ACVR1•ActA•IIR NSCs inhibit BMP signaling in HEK293[myc-ACVR1] and W20 cells. **a**,**b**, HEK293 cells overexpressing myc-human ACVR1 were either left untreated or treated with 10nM ActA for 1 hour. After washing off the unbound ActA, the cells were exposed to varying concentrations of BMP7 for either 30 minutes or 2 hours. Immunoblot analysis (**a**) and quantification (**b**) demonstrate that ActA significantly decreases BMP7-induced Smad1/5 phosphorylation after 30 minutes. BMP7-induced Smad1/5 phosphorylation recovered 2 hours after the ActA treatment. **c**, Expression of BMP/TGFß type I and type II receptors in W20 cells and Bmpr1a^−/-^; Bmpr1b^−/-^ mESCs. Type I receptor mRNA levels of W20 cells and Bmpr1a^−/-^; Bmpr1b^−/-^ mESCs were analyzed. W20 cells express 10-fold more ACVR1 mRNA and 8-fold more TGFBR1 mRNA than Bmpr1a^−/-^; Bmpr1b^−/-^ mESCs. The mRNA levels of ACVR1B in W20 cells is similar in both lines. **d**,**e**, W20 cells were either left untreated or treated with 10nM ActA for 1 hour. After washing off the unbound ActA, the cells were exposed to varying concentrations of BMP7 for either 10 or 30 minutes. Immunoblot analysis (**d**) and quantification (**e**) show ActA treatment significantly reduces the BMP7-induced Smad1/5 phosphorylation.

**Extended Data Figure 8.**
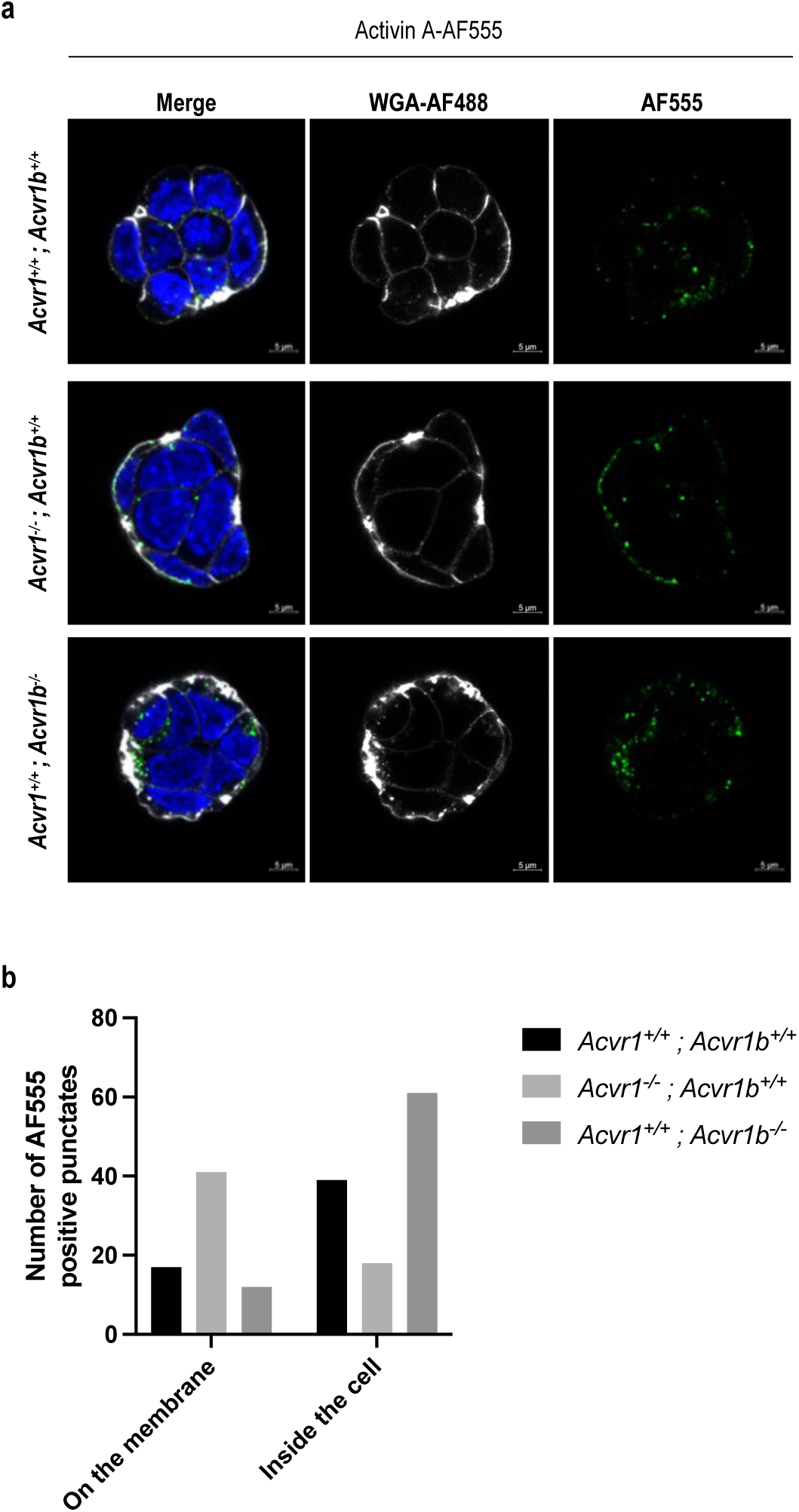
ACVR1B is not required for endocytosis of ActA. Wild type (Acvr1^+/+^; Acvr1b^+/+^), Acvr1 homozygous-null (Acvr1^−/-^; Acvr1b^+/+^), and Acvr1b homozygous-null (Acvr1^+/+^; Acvr1b^−/-^) mESCs were either treated with 30nM AF555-labeled ActA. **a**, Cells were stained with WGA-AF488 to mark the membrane, depicted in white, while AF555 and nuclei staining are represented in green and blue, respectively. Confocal microscopy was used for imaging. ActA-AF555 was not internalized in Acvr1 homozygous-null mESCs but was internalized in Acvr1b homozygous-null mESCs. **b**, Quantification of internalized and membrane-localized ActA-AF555 was done using Image J.

**Extended Data Figure 9.**
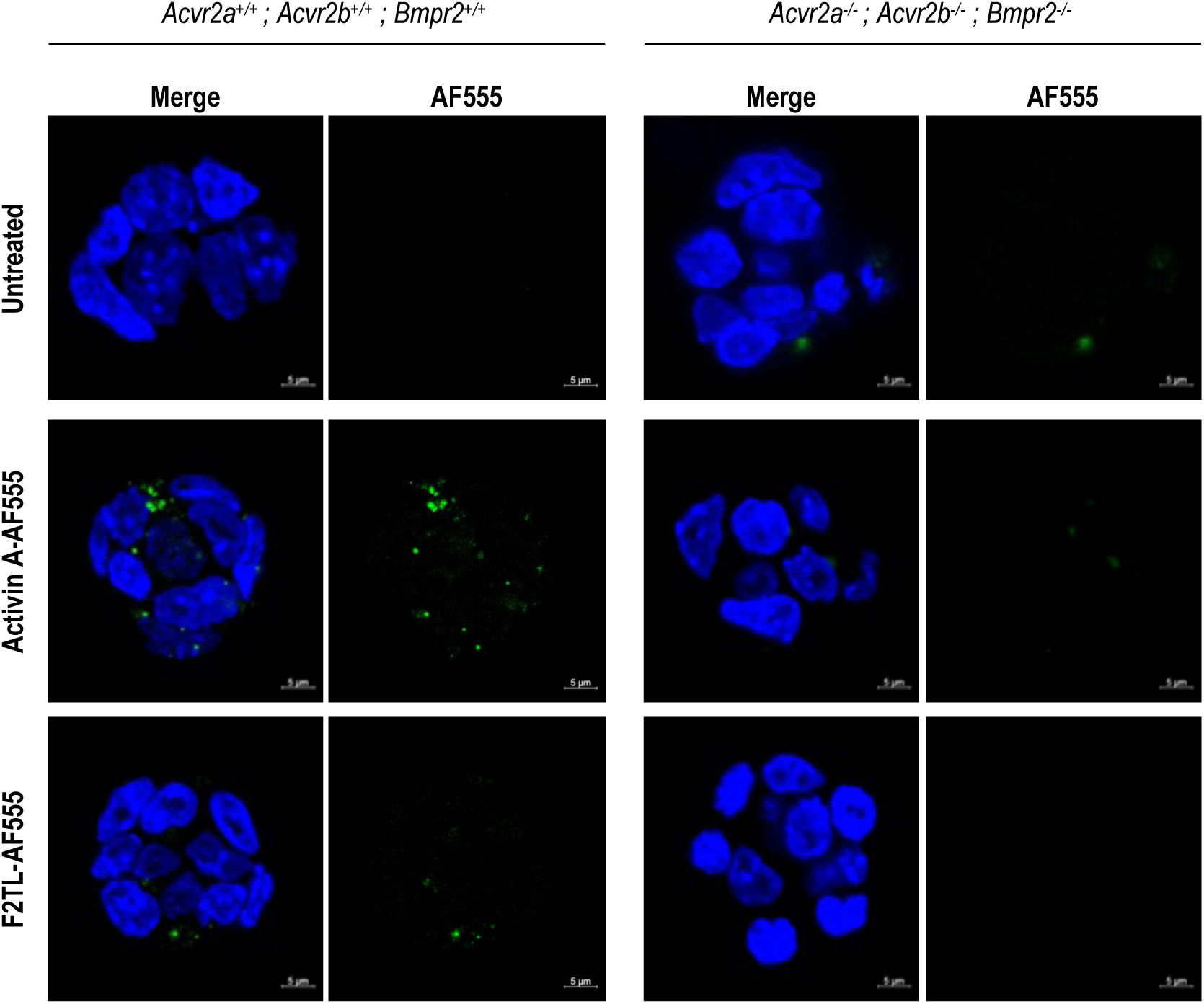
Type II receptors are required for endocytosis of ACVR1•ActA complexes. Acvr2a^+/ +^; Acvr2b^+/+^; Bmpr2^+/+^ mESCs and Acvr2a^−/-^; Acvr2b^−/-^; Bmpr2^−/-^ mESCs were treated with 30nM AF555-labeled ActA or F2TL or left untreated. Activin A-AF555 did not internalize in mESCs lacking the type II receptors utilized by ActA (Acvr2a^−/-^; Acvr2b^−/-^; Bmpr2^−/-^ mESCs) but slightly internalized in mESCs expressing endogenous levels of type II receptors. F2TL-AF555 did not internalize in either condition.

**Extended Data Figure 10.**
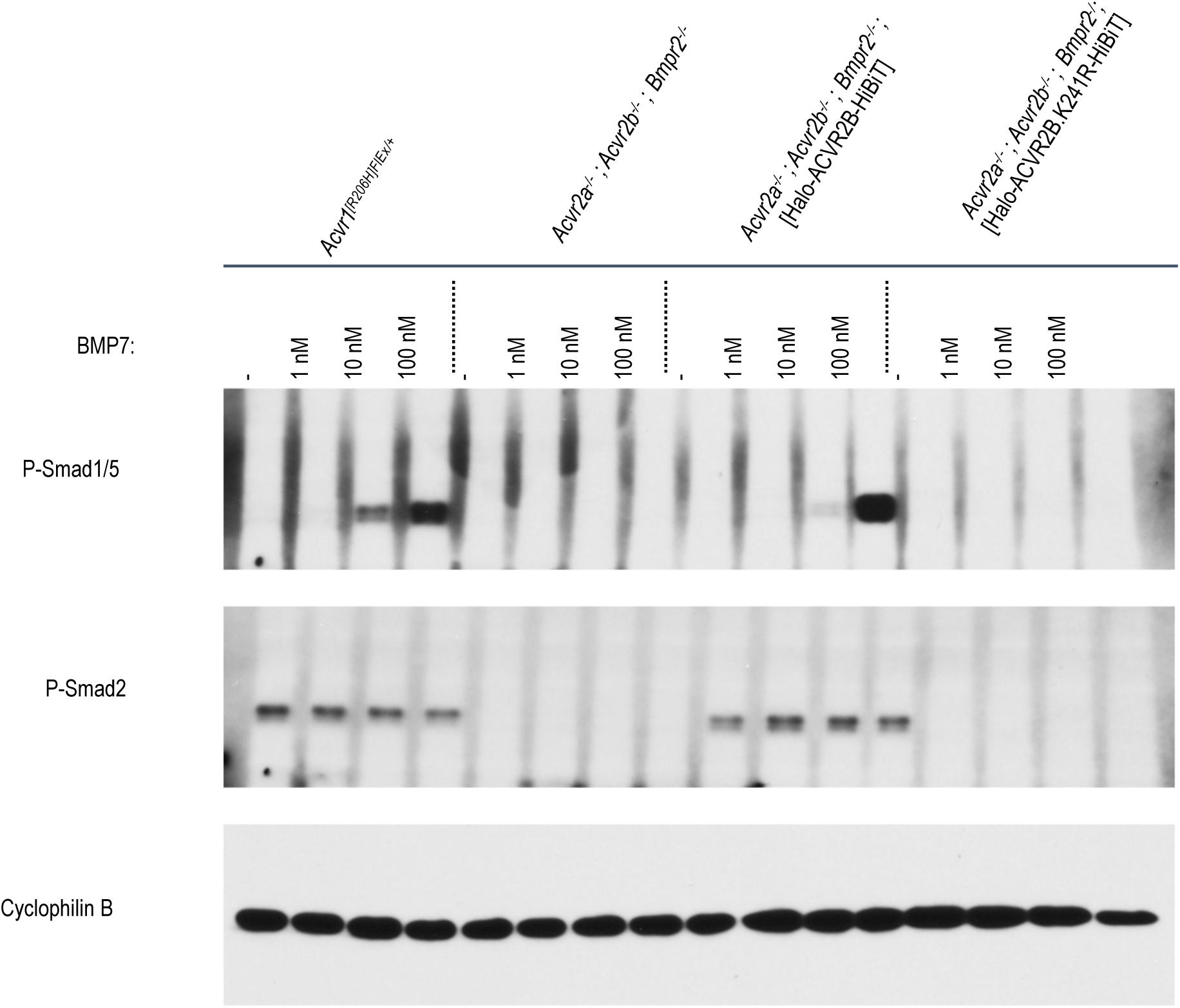
BMP7 induces Smad1/5 phosphorylation in Halo-ACVR2B-HiBiT-expressing mESCs. *Acvr1^[R206H]FlEx/+^* mESCs, *Acvr2a^−/-^; Acvr2b^−/-^; Bmpr2^−/--^*mESCs, and *Acvr2a^−/-^; Acvr2b^−/-^; Bmpr2^−/-^* mESCs expressing Halo-ACVR2B-HiBiT and Halo-ACVR2B.K241R-HiBiT were treated with varying concentrations of BMP7 for 1 hour. BMP7 induced Smad1/5 phosphorylation both in the parental *Acvr1^[R206H]FlEx/+^* and Halo-ACVR2B-HiBiT expressing mESCs. It did not induce Smad1/5 phosphorylation in the kinase-dead Halo-ACVR2B.K241R-HiBiT expressing mESCs.

## Online methods

### Reagents

ActA (338-AC-500/CF), Activin B (659-AB/CF), Activin AB (1066-AB-005/CF), Activin AC (4879-AC-010/CF), BMP7 (354-BP-010/CF), and anti-ALK4 antibody (MAB222) were purchased from R&D Systems. Activin A.Nod.F2TL (F2TL), anti-ACVR1 antibody (REGN5168), and hIgG4 isotype control antibody were expressed and purified in-house. The Activin A-AF555 and F2TL-AF555 were generated in-house.

### Generation and culturing of genetically modified mESCs

*Acvr1^−/-^*, *Acvr1b^−/-^*, *Bmpr1a^−/-^; Bmpr1b^−/-^*, and *Acvr2a^−/-^; Acvr2b^−/-^; Bmpr2^−/-^* mES cell lines were generated as previously described using CRISPR guides to biallelically ablate the target gene ^4^. mESC lines were expanded on gelatin-coated flasks, layered with irradiated primary mouse embryonic fibroblast (MEF) feeder cells from the DR4 mouse strain, which is resistant to neomycin, hygromycin, puromycin, and 6-thioguanine (MilliporeSigma, PMEF-DR4X). mESCs were cultured in VG-DMEM media containing 15% (v/v) FBS, 50 U/mL penicillin/streptomycin, 0.2% (v/v) b-mercaptoethanol, and 2U/mL leukemia inhibitory factor (MilliporeSigma, ESG1107) in a 37°C humidified incubator with 5% CO_2_. MEF feeder cells were removed using magnetic feeder removal microbeads (Miltenyi Biotec, 130-095-531) according to the manufacturer’s protocol. After the removal of feeder cells, the mESCs were cultured in 2i media ^54^ on gelatin-coated plates.

### Lentivirus and transduction of mESCs

Wild-type, mutant versions of HaloTag-mouse ACVR1-HiBiT, HaloTag-mouse ACVR1B-HiBiT and HaloTag-mouse ACVR2B-HiBiT lentiviruses were generated in HEK293T cells using Lenti-X Packaging Single Shots (Takara Bio, 631276) and concentrated 20-fold with Lenti Concentrator (OriGene, TR30025), in accordance with the manufacturers’ instructions. A mixture of 50µL of concentrated lentivirus and 20µL LentiBlast Premium transduction enhancer (Oz Biosciences, LBPX500) was combined with 1 × 10^6^ mESCs immediately after MEF feeder cell removal and then plated in gelatin-coated 6-well plates. After 48 hours, mESCs were transferred to gelatin coated T75 flasks containing MEF feeder cells to undergo antibiotic selection until stable cell lines were generated.

### Flow cytometry

mESCs were non-enzymatically dissociated using Accutase solution (MilliporeSigma, A6964) and resuspended in flow cytometry staining buffer (Thermo Fisher Scientific, 50-112-9748) after the removal of feeder cells. Following a 15-minute blocking period (Thermo Fisher Scientific, 14-9161-73), cells were stained for one hour with 3.5µM HaloTag Alexa Fluor 660 ligand. This was followed by a viability staining (Thermo Fisher Scientific, L34955) for 30 minutes at a 1:1000 dilution. The stained cells were then fixed with CytoFix (BD Biosciences, 554655), passed through a filter block (Pall, PN 8027), and analyzed using a CytoFLEX LX flow cytometer (Beckman Coulter).

### Confocal Microscopy

mESCs were plated in gelatin-coated 8-well chamber slides (Ibidi, 80826) in 2i media following the removal of feeder cells. After 48 hours, the 2i media was replaced with serum-free KnockOut DMEM (Thermo Fisher Scientific, 10829018) supplemented with 2mM L-glutamine, 50U/mL penicillin/streptomycin, 2 U/mL Leukemia Inhibitory Factor, and 0.1% bovine serum albumin (BSA) for 1 hour. During this incubation period, HaloTag-receptor expressing mESCs were fluorescently labeled with either 1 µM of HaloTag AlexaFluor 488 ligand (Promega, G1001) or 3.5µM of HaloTag AlexaFluor 660 ligand (Promega, G8471). After receptor labeling, media was replaced with fresh serum-free media containing 30nM fluorescently labeled ligands and the cells were incubated for 1 hour. The cell membrane of mESCs without HaloTag-receptors was fluorescently stained with wheat germ agglutinin Alexa Fluor 488 (Thermo Fisher Scientific, W11261) according to the manufacturer’s protocol. Following the 1-hour ligand treatment, cells were fixed with 4% paraformaldehyde (Biotium, 22023). For lysosomal staining, cells were permeabilized with ice-cold methanol at -20°C for 10 minutes, blocked in 10% BSA in PBS for 1 hour at room temperature, and stained with FITC anti-LAMP1 antibody (Abcam, ab24871) at 1:200 dilution overnight at 4°C. After fixation or after LAMP1 staining, ProLong Diamond Antifade Mountant with DAPI (Thermo Fisher Scientific, P36971) was added to stain cell nuclei. The cells were then imaged under a confocal microscope at 100x magnification (Zeiss, LSM710).

### Live cell imaging

HEK293 cells expressing human ACVR1-GFP were stained with Lysopainter (Abcam, ab176829) and Hoechst (ThermoScientific, 62249). Live cell imaging was done using Zeiss LSM880 confocal microscope equipped with a humidified and temperature-controlled incubation chamber. The GFP signal was bleached, followed by the addition of 30nM Activin A. Internalization of ACVR1-GFP was then monitored for a duration of 30 minutes. Co-localization of ACVR1-GFP and Lysopainter was calculated using the ZenBlue software.

### Cell lines, culture conditions, and transfections

HEK293 and W20 cell lines were cultured in DMEM, supplemented with 10% (v/v) FBS, 50U/mL penicillin/streptomycin, and 2mM L-glutamine in a humidified 37°C incubator with 5% CO_2_. Transfection of the HEK293 cells was carried out using the TransIT-293 DNA transfection reagent (MirusBio, MIR 2700), following the manufacturer’s recommended protocol.

### Immunoprecipitation

The HEK293 cells were transfected with HA-human ACVR2B (HA-ACVR2B) and either myc-human ACVR1 (myc-ACVR1) or myc-human ACVR1.R206H (myc-ACVR1.R206H). After a period of 48 hours, the cells were switched to serum-free DMEM for 1 hour, followed by a treatment with 10nM of ligand for an additional hour. Membrane fractions were then collected using the Mem-PER Plus membrane protein extraction kit (Thermo Fisher Scientific, 89842), in accordance with the manufacturer’s instructions. The protein concentration of the membrane fractions and whole cell lysates were measured using the Pierce BCA protein assay kit (Thermo Fisher Scientific, 23227). Myc-tag immunoprecipitation (IP) was carried out using the Myc-tag magnetic IP kit (Thermo Fisher Scientific, 88844), following the manufacturer’s recommended protocol.

### Ligand treatments and immunoblotting

Forty-eight hours after plating cells, media was switched to serum-free media for 1 hour followed by 1 hour treatment with various ligands or antibodies. Whole-cell lysates were collected with RIPA lysis buffer (Thermo Fisher Scientific, 89901) containing 2X protease and phosphatase inhibitor (Thermo Fisher Scientific, 78440) and 0.1U/mL nuclease (Thermo Fisher Scientific, 88701). Equal amounts of lysates were resolved under reducing conditions in 4%-20% Novex WedgeWell gels (Invitrogen) and transferred to PVDF membranes (Advansta, L-08004-010). Membranes were blocked for 2 hours at room temperature with SuperBlock T20 blocking buffer (Thermo Fisher Scientific, 37536) and incubated with primary antibodies at a 1:1000 dilution (anti-P-Smad1/5 [clone 41D10], anti-P-Smad2 [clone 138D4]), 1:4000 dilution (anti-Myc [clone 71D10], anti-HA [clone C29F4]), 1:10,000 dilution (anti-sodium potassium ATPase [clone EP1845Y]-plasma membrane loading control) or 1:10,000 dilution (anti-cyclophilin B [clone D1V5J]-equal protein loading control) overnight at 4°C. Membranes were then incubated with horseradish peroxidase-conjugated secondary antibody at a range of dilutions from 1:2000 – 1:10,000 (depending on the primary antibody) for 2 hours at room temperature. Western Bright ECL HRP substrate was used for detection (Advansta, K-12045-D20). Band intensities of the western images were analyzed using ImageJ.

### Luciferase reporter assay

Stable pools of HEK293-BRE-luciferase (Smad1/5-responsive) and HEK293-CAGA-luciferase (Smad2/3 responsive) reporter cells were generated. Approximately 8,000 cells were plated in a 96-well plate in complete media. Media was replaced with serum-free DMEM, supplemented with 2mM L-glutamine, 50U/mL penicillin/streptomycin, and 0.1% BSA, twenty-four hours post-plating. Following an overnight starvation period, ligands at various concentrations were added. After 16 hours, luciferase expression was measured using the Bright-Glo luciferase assay system (Promega, E2650).

